# Elevated NAD^+^ drives Sir2A-mediated GCβ deacetylation and OES localization for *Plasmodium* ookinete gliding and mosquito infection

**DOI:** 10.1101/2024.10.03.616407

**Authors:** Yang Shi, Lin Wan, Mengmeng Jiao, Chuan-qi Zhong, Huiting Cui, Jing Yuan

## Abstract

cGMP signal-activated ookinete gliding is essential for mosquito midgut infection of *Plasmodium* in malaria transmission. During ookinete development, cGMP synthesizer GCβ polarizes to a unique localization <ookinete extrados site= (OES) until ookinete maturation and activates cGMP signaling for initiating parasite motility. However, the mechanism underlying GCβ translocation from cytosol to OES remains elusive. Here, we used protein proximity labeling to search the GCβ-interacting proteins in ookinetes of the rodent malaria parasite *P. yoelii*, and found the top hit Sir2A, a NAD^+^-dependent sirtuin family deacetylase. Sir2A interacts with GCβ throughout ookinete development. In mature ookinetes, Sir2A co-localizes with GCβ at OES in a mutually dependent manner. Parasites lacking Sir2A lose GCβ localization at OES, ookinete gliding, and mosquito infection, phenocopying GCβ deficiency. GCβ is acetylated at gametocytes but is deacetylated by Sir2A for OES localization at mature ookinetes. We further demonstrated that the level of NAD^+^, an essential co-substrate for sirtuin, increases during the ookinete development. The NAD^+^ at its maximal level until ookinete maturation promotes Sir2A-catalyzed GCβ deacetylation, ensuring GCβ localization at OES. This study highlights the spatiotemporal coordination of cytosolic NAD^+^ level and NAD^+^-dependent Sir2A in regulating GCβ deacetylation and dynamic localization for *Plasmodium* ookinete gliding.

## Introduction

Malaria, caused by the protozoan parasite *Plasmodium*, is an infectious disease resulting in an estimated 49 million cases and 608,000 deaths globally in 2022^1^. The spread of malaria relies on parasite infection and development in the mosquito vector. Once entering the mosquito midgut after a blood meal, male and female gametocytes are immediately activated into male and female gametes, which fertilize to form the zygotes. Within 12 to 20 hours, the spherical zygotes undergo remarkable morphogenesis of <protrusion-elongation-maturation= to differentiate into crescent-shaped ookinetes^2,3^. Only mature ookinetes activating gliding motility could move through the blood bolus and traverse the midgut epithelium barrier. Following colonization at the midgut basal lamina, the ookinete develops into an oocyst, each giving rise to thousands of sporozoites^4,5^. When mosquitoes bite again, the sporozoites in the salivary glands are injected into another vertebrate host.

Ookinete motility is powered by the glideosome, an actomyosin-based protein machinery located between the parasite plasma membrane (PPM) and the underneath membranal organelle of the inner membrane complex (IMC)^6,7^. The mechanical force produced by glideosome is converted to the backward movement of the adhesin protein CTRP^8^, generating parasite forward movement for gliding or invasion^9^. 39-59-cyclic guanosine monophosphate (cGMP), guanylate cyclase beta (GCβ), phosphodiesterase delta (PDE·), and cGMP-dependent protein kinase G (PKG) play critical roles in upstream signaling of ookinete motility. In mature ookinete, the gliding initiation depends on the activation of cGMP signal^10–12^, which level is coordinated by the activities of GCβ (synthesizes cGMP) and PDE· (hydrolyzes cGMP)^12^. Only the cGMP level exceeding the threshold could activate PKG, resulting in PLC/IP3-mediated Ca^2+^ release, phosphorylation of multiple glideosome proteins, and initiation of ookinete gliding^11–13^.

*Plasmodium* parasites encode two guanylyl cyclases GCβ and GCβ^14–16^. Both GCs are large proteins (3000-4000 amino acids in length) possessing an unusual protein architecture, in which the C-terminal guanylate cyclase domain (GCD) is combined with an unrelated N-terminal P4-type ATPase-like domain (ALD)^17,18^. While the GCD is responsible for cGMP synthesis, the function of the ALD is still obscure^14–16^. GCβ is genetically essential for ookinete gliding in *P. berghei*, *P yoelii*, and *P. falciparum*^10–12,19^. Due to the large size, multiple transmembrane helixes, and the ALD-GCD hybrid domain structure of GCβ, the geography of GCβ-mediated cGMP signaling in the *Plasmodium* ookinete gliding remained elusive for a long time^20–22^.

As our previous efforts to investigate the expression and localization of GCβ and PDE· in ookinete gliding, we revealed a spatiotemporal regulation of cGMP signal in the *P. yoelii*^19^. During the ookinete development, GCβ and PDE· are distributed in the cytoplasm. Until ookinete maturation, GCβ translocates and polarizes to PPM at the <ookinete extrados site= (OES) while PDE· maintains cytosolic^19^. The OES is a subapical area in the outer curve of the crescent ookinete^19,21,22^. GCβ polarization at OES initiates the gliding of mature ookinete. In addition, the P4-ATPase co-factor CDC50A is also localized at OES and functions as a chaperone to stabilize GCβ^19^. Based on these results, we proposed a GCβ/CDC50A polarization-directed cGMP signal activation model for ookinete gliding^19^. Before ookinete maturation, GCβ/CDC50A complex and PDE· maintain a sub-threshold cGMP level precluding PKG activation in the cytoplasm. Upon ookinete maturation, GCβ/CDC50A complex translocates to OES. The GCβ/CDC50A polarization increases the local cGMP concentration that drives PKG activation and initiates ookinetes gliding^19^. Despite the progress in understanding cGMP signaling of ookinete gliding, the mechanism underlying GCβ translocation from cytosol to OES until mature ookinete remains unknown.

Protein acetylation is a posttranslational modification regulating protein stability, localization, and protein-protein interaction^23,24^. Protein acetylation is dynamically controlled by acetylase and deacetylase^24,25^. The silent information regulator 2 family proteins of deacetylase (Sirtuin or SIRT) are found in organisms ranging from bacteria to humans^26^. Sirtuins deacetylate the acetyl-lysine residue from the acetylated substrates and its catalytic activity depends on the level of the nicotinamide adenine dinucleotide (NAD^+^)^27,28^. *Plasmodium* parasites encode two sirtuin proteins Sir2A and Sir2B^29^. The *P. falciparum* parasite evolves the antigenic variation at the infected erythrocyte surface to avoid host immune clearance via regulated gene expression of the *var* gene family-encoded antigens^30,31^. In the asexual blood stages of *P. falciparum*, Sir2A is thought to regulate histone deacetylation in the telomeric regions, and control the antigenic expression of the *var* genes^30^. So far, the role of Sir2A has not been investigated in the mosquito stages of *Plasmodium*.

In this study, we searched the GCβ-interacting proteins at OES in the *P. yoelii* ookinetes and found that Sir2A forms a complex with GCβ/CDC50A during ookinete development. Parasites lacking Sir2A phenocopy GCβ deficiency in ookinete gliding. Sir2A catalyzes GCβ deacetylation and modulates GCβ localization at OES in mature ookinete. We further demonstrated that the level of NAD^+^, an essential co-substrate of Sir2A, increases during the ookinete development. The NAD^+^ at its maximal level promotes Sir2A-catalyzed GCβ deacetylation until ookinete maturation.

## Results

### Proximity proteomics identifies Sir2A a potential GCβ-interacting protein in ookinete of *P. yoelii*

To search potential regulators of GCβ localization at OES, we applied the biotin ligase TurboID-based proximity labeling (PL) to track the GCβ-interacting proteins in the ookinetes. The endogenous GCβ was fused with a TurboID::6HA domain via CRISPR-Cas9 in the *P. yoelii* 17XNL strain, generating the modified line *gcβ::TurboID* (*Tb-GCβ* in short) (Figure 1A). We generated a control line *gcβ::T2A::TurboID::6HA* (*Tb-Cyto* in short), in which a T2A peptide was inserted to direct cytosolic expression of TurboID::6HA alone under the promoter of *gcβ* gene (Figure 1A). As expected, the fusion protein Tb-GCβ was localized at OES while the Tb-Cyto was cytosolic in the ookinetes (Figure 1A). After incubation with 50 µM biotin for 3 hours at 22°C, the ookinetes expressing ligase were co-stained with fluorescence-conjugated streptavidin and anti-HA antibody. As expected, the biotinylated proteins were co-localized with the fusion ligase at OES in the *Tb-GCβ* ookinetes, while the biotinylated proteins and the ligase were in cytosolic in *Tb-Cyto* ookinetes (Figure 1A). Three biological replicates were prepared from the *Tb-GCβ* and *Tb-Cyto* ookinetes, and the streptavidin-affinity purified proteins from cell extracts were subjected to proteomic analysis. Quantitative mass spectrometry yielded 251 enriched proteins with high confidence in *Tb-GCβ* compared to *Tb-Cyto* ookinetes (Figure 1B, Supplementary Data 1). GCβ, after being cis-biotinylated, was included in these protein hits (blue dot in Figure 1B). CDC50A, an essential cofactor of GCβ^19^, was also detected (blue dot in Figure 1B), indicating the good quality of these PL experiments. Among the significant hits, the top is the sirtuin family protein Sir2A (red dot in Figure 1B). So far, the expression, localization, and function of Sir2A have not been investigated in mosquito stages of *Plasmodium*, including the ookinete.

**Figure 1.**
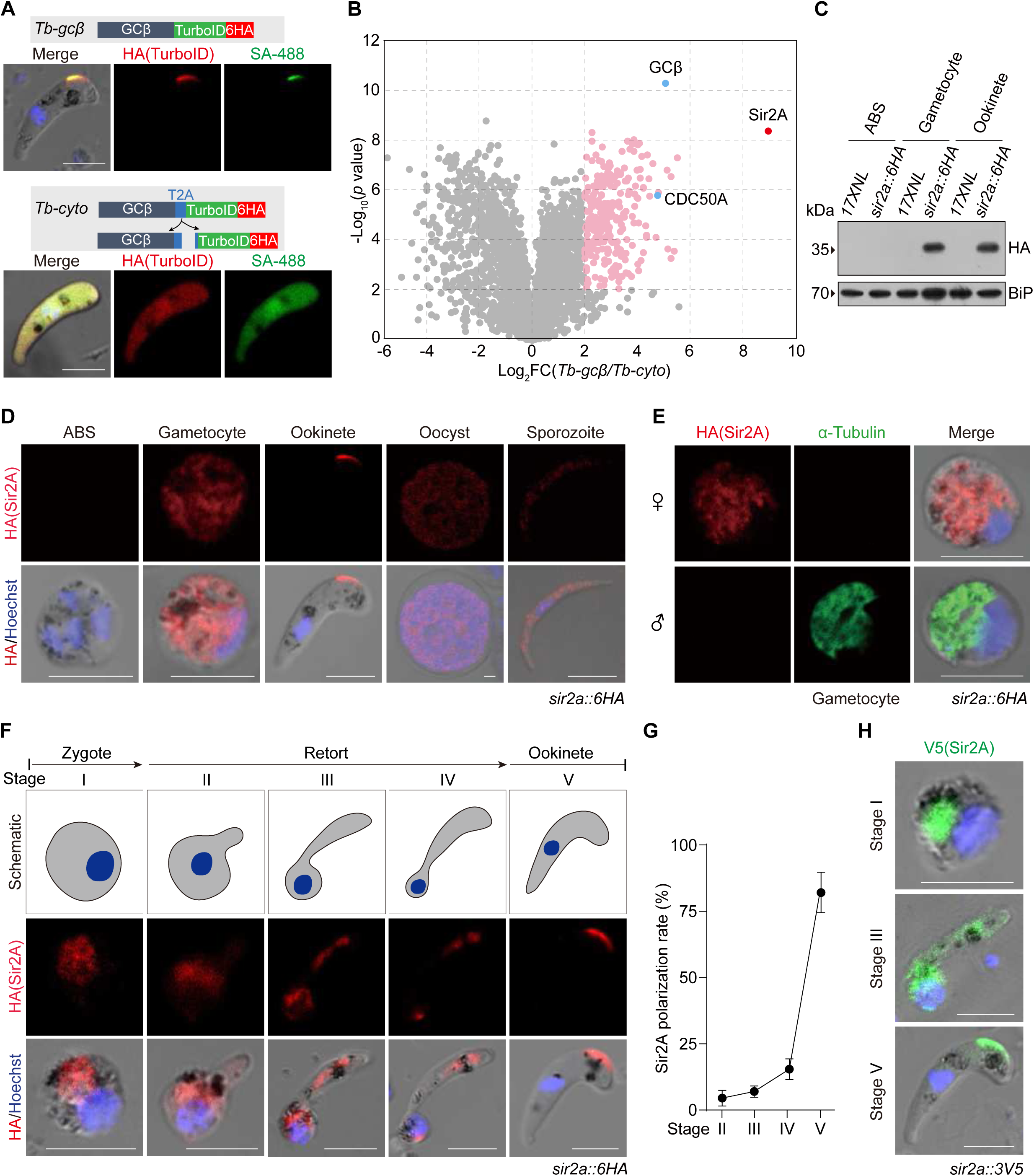
Proximity proteomics identifies Sir2A a potential GCβ-interacting protein in ookinete of *P. yoelii*. **A.** Schematic of the modified line used for TurboID ligase-mediated proximity labeling of GCβ-interacting proteins in living ookinetes. Endogenous GCβ was C-terminally tagged with a TurboID::6HA motif by CRISPR-Cas9 in 17XNL, generating *Tb-gcβ* line. *Tb-Cyto* is a control line in which the T2A is inserted between GCβ and TurboID for separated expression of GCβ and TurboID. Co-staining of HA-tagged TurboID ligase (red) and biotinylated proteins (SA-488, green) in ookinetes after incubated with 50 µM biotin at 22 °C for 3 h. Scale bars: 5 μm. Three independent experiments. **B.** Volcano plots showing 251 significantly enriched proteins (cutoffs log_2_FC)g)1 and *p*-value f 0.05) in *Tb-gcβ* versus *Tb-Cyto* ookinetes. Protein list is in Supplementary Data 1. The relative enrichment ratio (x axis) of protein was calculated by quantifying protein intensity in *Tb-gcβ* relative to *Tb-Cyto* (n=3). The *p*-values were calculated by two-sided *t*-test and adjusted by FDR. GCβ, CDC50A, and Sir2A are highlighted. **C.** Immunoblot of Sir2A in asexual blood stages (ABS), gametocytes, and ookinetes of 17XNL and *sir2a::6HA* parasites. BiP was used as a loading control. Three independent experiments. **D.** Immunofluorescence assay (IFA) detecting Sir2A expression in multiple stages of the *sir2a::6HA* parasite. The parasites were co-stained with anti-HA antibody and DNA dye Hoechst 33342. Scale bar: 5 μm. Three independent experiments. **E.** Co-staining *sir2a::6HA* gametocytes with antibodies against HA and α-Tubulin II (male gametocyte specific protein). Scale bar: 5 μm. Three independent experiments. **F.** IFA of Sir2A expression dynamics during the zygote to ookinete development of *sir2a::6HA* parasite. Scale bars: 5 μm. Three independent experiments. **G.** Quantification of Sir2A polarization level at OES during the *in vitro* ookinete development in **F**. Polarization rates are mean ± SD of three independent experiments. Thirty ookinetes were analyzed for each group in each experiment. **H.** IFA of Sir2A expression dynamics during the zygote to ookinete development of *sir2a::3V5* parasite. Scale bars: 5 μm. Three independent experiments.

To validate the PL results and investigate the expression and localization of Sir2A during the parasite life cycle of *P. yoelii*, we tagged the endogenous Sir2A (PY17X_1348600) with a sextuple HA epitope (6HA) at C-terminus in the 17XNL strain using CRISPR/Cas9^32,33^. The tagged parasite line *sir2a::6HA* developed normally in mice and mosquitoes, indicating no detectable detrimental effect of tagging on protein function. Immunoblot and immunofluorescence assay (IFA) showed that Sir2A was expressed in gametocytes, ookinetes, mosquito midgut oocysts, and mosquito salivary gland sporozoites, but was not detected in asexual blood stage parasites (Figure 1C and D). Co-staining of the *sir2a::6HA* gametocytes with β-Tubulin (male gametocyte highly expressed) and HA antibodies showed that Sir2A was expressed only in female gametocytes (Figure 1E). Interestingly, Sir2A was distributed in the cytoplasm of gametocytes, oocysts, and sporozoites, but was concentrated at a site posterior to the apical of ookinetes (Figure 1D). This area in ookinetes was designated as the 88ookinete extrados site99 (OES)^19^. During the *in vitro* zygote to ookinete differentiation, Sir2A was distributed in the cytoplasm from zygote to retort, but concentrated to OES in mature ookinetes (Figure 1F). We quantified the polarization level of Sir2A by calculating fluorescent signals at OES over the whole cell at different stages of ookinete development (Figure G). We generated another parasite line *sir2a::3V5* in which the endogenous Sir2A was tagged with a triple V5 epitope (3V5) in the C-terminus and observed similar expression and localization of Sir2A during zygote to ookinete differentiation (Figure 1H). Together, we identified Sir2A as a potential GCβ-interacting protein at OES in the ookinetes. Sir2A had a dynamical localization pattern during zygote to ookinete development, similar to the GCβ/CDC50A complex^19^.

### Sir2A co-localizes and interacts with GCβ/CDC50A during ookinete development

We investigated the temporal-spatial association between Sir2A and GCβ during the gametocyte-zygote-ookinete development. From the parasite *gcβ::6HA* generated previously^19^, the endogenous Sir2A was tagged with a 3V5 at the C-terminus, generating a double-tagged line *gcβ::6HA*;*sir2a::3V5*. In this line, we analyzed the time-course localization dynamic of Sir2A and GCβ. In the female gametocytes, Sir2A and GCβ were cytosolic but not overlaid with each other (Figure 2A). After gamete fertilization, Sir2A became co-localized with GCβ at the cytosol in the zygotes until later-stage retorts. In mature ookinetes, both Sir2A and GCβ were concentrated to OES (Figure 2A). We used different methods to analyze the association between Sir2A and GCβ during gametocyte-zygote-ookinete development. Co-immunoprecipitation (Co-IP) also revealed that Sir2A did not interact with GCβ in gametocytes of the *gcβ::6HA*;*sir2a::3V5* (*DTS1*) parasites (Figure 2B). As a positive control, GCβ bound its cofactor CDC50A in gametocytes of the *gcβ::6HA*;*50a::3V5* (*DTS2*) parasites^19^ (Figure 2B). Proximity Ligation Assay (PLA) is an immunohistochemical tool to detect protein interaction with specificity and sensitivity^34^. As a positive control, PLA signals indicating GCβ and CDC50A interaction were detected in the *gcβ::6HA*;*50a::3V5* gametocytes (Figure 2C). No PLA signal was detected in the *gcβ::6HA*;*sir2a::3V5* gametocytes when using both anti-HA and anti-V5 antibodies (Figure 2C). In contrast, both Co-IP and PLA detected that Sir2A co-localized and interacted with GCβ at cytosol in the zygotes (Figure 2D and E) and at OES in the ookinetes (Figure 2F and G).

**Figure 2.**
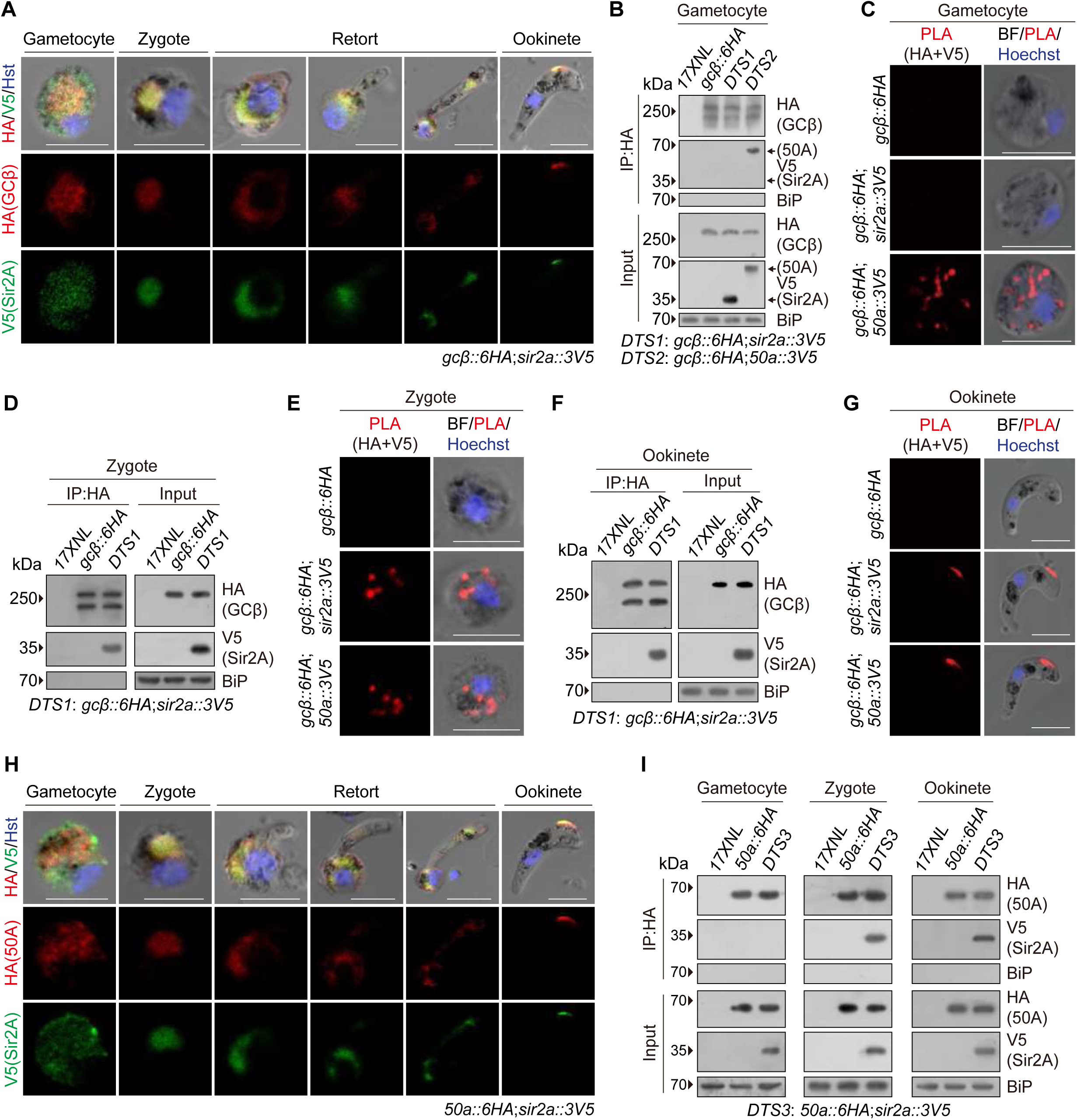
Sir2A co-localizes and interacts with GCβ/CDC50A during ookinete development. **A.** IFA of HA-tagged GCβ and V5-tagged Sir2A expression during the gametocyte to ookinete development of the *gcβ::6HA*;*sir2a::3V5* parasite. Scale bars: 5 μm. Three independent experiments. **B.** Co-immunoprecipitation (Co-IP) of GCβ and Sir2A in gametocytes of the *gcβ::6HA*;*sir2a::3V5* (*DTS1*) parasite. Co-IP was conducted using anti-HA antibody. BiP as a loading control. Interaction between GCβ and CDC50A in *gcβ::6HA*;*cdc50a::3V5* (*DTS2*) gametocytes was used as a positive control. Two independent experiments. **C.** Proximity ligation assay (PLA) detecting protein interaction between GCβ and Sir2A in the *gcβ::6HA*;*sir2a::3V5* gametocytes. GCβ and CDC50A interaction in *gcβ::6HA*;*cdc50a::3V5* gametocytes was used as a positive control. Scale bars: 5 μm. Two independent experiments. **D.** Co-IP of GCβ and Sir2A in the *DTS1* zygotes. Co-IP was conducted using anti-HA antibody. BiP as a loading control. Two independent experiments. **E.** PLA detecting protein interaction between GCβ and Sir2A in the *DTS1* zygotes. Scale bars: 5 μm. Two independent experiments. **F.** Co-IP of GCβ and Sir2A in the *DTS1* ookinetes. Co-IP was conducted using anti-HA antibody. BiP as a loading control. Two independent experiments. **G.** PLA detecting protein interaction between GCβ and Sir2A in the *DTS1* ookinetes. Two independently performed experiments with similar results. Scale bars: 5 μm. **H.** IFA of HA-tagged CDC50A and V5-tagged Sir2A expression during the gametocyte to ookinete development of the *50a::6HA*;*sir2a::3V5* parasite. Scale bars: 5 μm. Three independent experiments. **I.** Co-IP of CDC50A and Sir2A in gametocyte, zygote, and ookinete of the *cdc50a::6HA*;*sir2a::3V5* (*DTS3*) parasite. Co-IP was conducted using anti-HA antibody. BiP as a loading control. Two independent experiments.

Since CDC50A is a cofactor of GCβ^19^, we also analyzed the association between Sir2A and CDC50A in this development. From the previous parasite line *cdc50a::6HA*^19^, we tagged the endogenous Sir2A with a 3V5 and obtained another double-tagged line *cdc50a::6HA*;*sir2a::3V5* (*DTS3*). Sir2A co-localized and interacted with CDC50A in zygotes and ookinetes in both IFA and Co-IP (Figure 2H and I), showing a similar association pattern as with GCβ. These results demonstrated that Sir2A does not interact with GCβ/CDC50A in the gametocytes but forms a complex with GCβ/CDC50A during the zygote to ookinete development.

### Sir2A phenocopies GCβ in regulating ookinete gliding for mosquito midgut infection

The *P. yoelii* Sir2A (PY17X_1348600) is a 278 amino acid protein that shows a high identity to Sir2A proteins from the rodent malaria parasite *P. berghei* and human malaria parasites *P. falciparum* and *P. vivax* (Supplementary Figure 1). To elucidate the function of Sir2A in the ookinetes, we deleted the whole coding region (837 bp) of the *sir2a* gene by homologous recombination via CRISPR-Cas9 in the *P. yoelii* 17XNL strain (wild-type) and obtained a mutant clone Δ*sir2a* (Figure 3A). The Δ*sir2a* exhibited normal asexual blood stage proliferation and gametocyte formation in the mice (Figure 3B and C). To evaluate the role of Sir2A in parasite development in the mosquito, *Anopheles stephensi* mosquitoes were fed on the parasite-infected mice. Δ*sir2a* produced no oocyst in the midgut on day 7 post-infection (pi) (Figure 3D) and no sporozoites in the salivary glands on day 14 pi (Figure 3E), indicating parasite transmission failure in the mosquito. As a parallel test, the GCβ-null parasite Δ*gcβ* failed to develop both oocyst and sporozoite in the mosquitoes as expected^12^ (Figure 3D and E).

**Figure 3.**
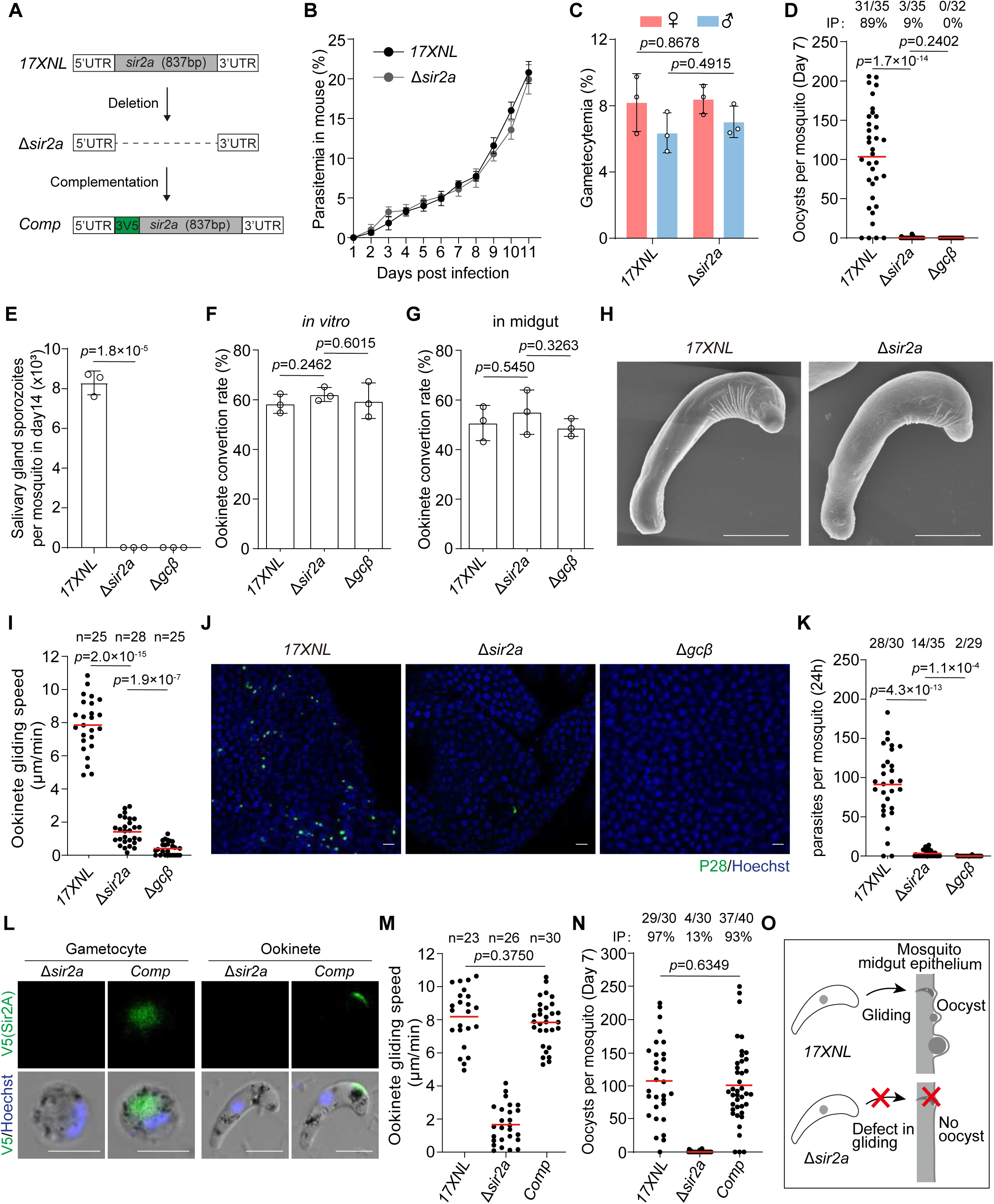
Sir2A phenocopies GCβ in regulating ookinete gliding for mosquito midgut infection. **A.** Schematic of the *sir2a* gene deletion and complementation using CRISPR-Cas9. The whole coding region of *sir2a* was removed in the 17XNL parasite, generating the Δ*sir2a* mutant. The *sir2a* gene from *P. yoelii* fused with a 3V5 was introduced back to the *sir2a* locus of the Δ*sir2a* mutant, generating complemented line *Comp*. **B.** Parasite proliferation at asexual blood stages in mice. Mean ± SD from three mice in each group. Two independent experiments. **C.** Male and female gametocyte formation in mice. Mean ± SD from three mice in each group, two-tailed *t*-test. Two independent experiments. **D.** Midgut oocyst formation in mosquitos at day 7 post infection. x/y on the top is the number of mosquitoes containing oocyst/the number of dissected mosquitoes, and the percentage represents the infection prevalence of mosquitoes. Red lines show the mean value. Two-sided Mann-Whitney *U* test. Two independent experiments. **E.** Salivary gland sporozoite formation in mosquitoes at day 14 post infection. Thirty infected mosquitoes were counted in each group. Mean ± SD from three independent experiments, two-tailed *t*-test. **F.** Mature ookinete formation *in vitro*. Mean ± SD from three independent experiments, two-tailed *t*-test. **G.** Mature ookinete formation in the mosquito midgut. Mean ± SD from three independent experiments, two-tailed *t*-test. **H.** Scanning electron microscopy (SEM) of 17XNL and Δ*sir2a* ookinetes. Scale bars: 5 μm. Three independent experiments. **I.** Ookinete gliding motility using the *in vitro* Matrigel-based assay. n is the number of ookinetes analyzed. Red lines show the mean value. Two-sided Mann-Whitney *U* test. Three independent experiments. **J.** IFA of P28 in ookinete and early oocyst at mosquito midguts infected with 17XNL, Δ*sir2a,* and Δ*gcβ* parasites 24 h post infection. P28 is a plasma membrane protein of ookinete and early oocyst. Scale bars: 20 μm. Three independent experiments. **K.** Quantification of parasites in **J**. x/y on the top is the number of midguts containing parasite/the number of midguts measured, red lines show the mean value. Two-sided Mann-Whitney test. **L.** IFA of the V5-tagged Sir2A in gametocytes and ookinetes of the complemented line *Comp*. Scale bar: 5 μm. Three independent experiments. **M.** Ookinete gliding motility. n is the number of ookinetes analyzed. Red lines show the mean value. Two-sided Mann-Whitney *U* test. Three independent experiments. **N.** Midgut oocyst formation in mosquitoes at day 7 post infection. Red horizontal lines show mean value, two-sided Mann-Whitney *U* test. x/y on the top is the number of mosquitoes containing oocyst/the number of mosquitoes dissected. The percentage represents the infection prevalence of mosquitoes. Two independent experiments. **O.** Cartoon showing Sir2A deficiency in ookinete gliding for mosquito midgut invasion.

The *Plasmodium* undergoes a gametocyte-gamete-zygote-ookinete development in the mosquito midgut and form the crescent-shaped motile ookinetes for penetration. We performed experiments to delineate the steps affected by Sir2A deficiency. The Δ*sir2a* showed normal male and female gamete formation *in vitro* compared with 17XNL (Supplementary Figure 2A and B). The *in vitro* assay for zygote to ookinete differentiation revealed that *sir2a* deletion had no marked effect on ookinete formation (Figure 3F). We isolated the Δ*sir2a* parasites from infected mosquito midguts and observed normal ookinetes (Figure 3G). Under scanning electron microscopy, the Δ*sir2a* mature ookinetes displayed a typical crescent shape as 17XNL (Figure 3H), suggesting no defect in the parasite morphology after loss of Sir2A. Since gliding motility is a prerequisite for the midgut traversal of ookinetes, we assessed the gliding capability of ookinetes using an *in vitro* Matrigel-based assay^11,35^. The Δ*sir2a* ookinetes displayed significantly reduced gliding speed compared to 17XNL but were slightly faster than the Δ*gcβ* ookinetes (17XNL: 7.9±1.6 μm/min, n=25; Δ*sir2a*: 1.4±0.8 μm/min, n=28; and Δ*gcβ*: 0.4±0.4 μm/min, n=25) (Figure 3I). The gliding-deficient ookinete of Δ*sir2a* may fail to traverse mosquito midgut. To test it, the midguts from infected mosquitoes were dissected at 24 hours pi (hpi) and visualized after staining with an antibody against P28 (a plasma membrane protein of ookinetes and early oocysts). Reduced numbers of P28^+^ parasites were detected in the midguts infected with Δ*sir2a* compared with 17XNL (midgut-associated parasites per mosquito: 91±45 in 17XNL, n=30; 3±4 in Δ*sir2a*, n=35; 0±0 in Δ*gcβ*, n=29) (Figure 3K).

To confirm that the ookinete gliding defect was caused by Sir2A deletion, we re-introduced the deleted 837 bp part fused with a 3V5 back into the *sir2a* locus of the Δ*sir2a* parasite (Figure 3A). Expression of the V5-tagged Sir2A was detected in gametocytes and ookinetes of the complemented line *Comp* (Figure 3L). In line with the endogenous Sir2A, the 3V5::Sir2A fusion protein exhibited OES localization in the *Comp* ookinetes (Figure 3L). Notably, the *Comp* parasite restored the oocyst and sporozoite formation in the infected mosquitoes (Figure 3M and N). Furthermore, we generated another Sir2A null parasite line by deleting the *sir2a* gene in the *sir2a::6HA* parasite (Supplementary Figure 2C and D). Consistent with the phenotypes for the Δ*sir2a* parasites, the mutant parasite *sir2a::6HA*;Δ*sir2a* displayed similar defects in the ookinete gliding and mosquito transmission of the parasite (Supplementary Figure 2E to K). These results demonstrated that Sir2A regulates ookinete gliding (Figure 3O), similar to the GCβ/CDC50A complex^19^.

### Mutual dependent localization of Sir2A and GCβ at OES in mature ookinete

We investigated whether Sir2A regulates GCβ expression or localization. The *sir2a* gene was deleted in the *gcβ::6HA* parasite, generating the mutant line *gcβ::6HA*;Δ*sir2a*. Immunoblot showed that Sir2A depletion did not affect protein levels of GCβ in gametocytes, zygotes, and ookinetes (Figure 4A), ruling out an effect of Sir2A on GCβ protein synthesis or stability. IFA found that Sir2A depletion did not affect the cytosolic distribution of GCβ in gametocytes and zygotes, but GCβ lost polarization at OES and was distributed at cytosol in the *gcβ::6HA*;Δ*sir2a* ookinetes (Figure 4B and C, Supplementary Figure 3). To further validate the localization alteration of GCβ in ookinetes after loss of Sir2A, we isolated the heavy fraction (including pellicle membrane and cytoskeleton) and light fraction (including cytoplasm) from the extracts of ookinete after hypotonic lysis. Immunoblot detected GCβ in heavy fraction from the *gcβ::6HA* ookinetes, but mainly in light fraction from the *gcβ::6HA*;Δ*sir2a* ookinetes (Figure 4D). These results in both IFA and protein fraction assays indicate that Sir2A is critical for GCβ localization at OES in ookinete.

**Figure 4.**
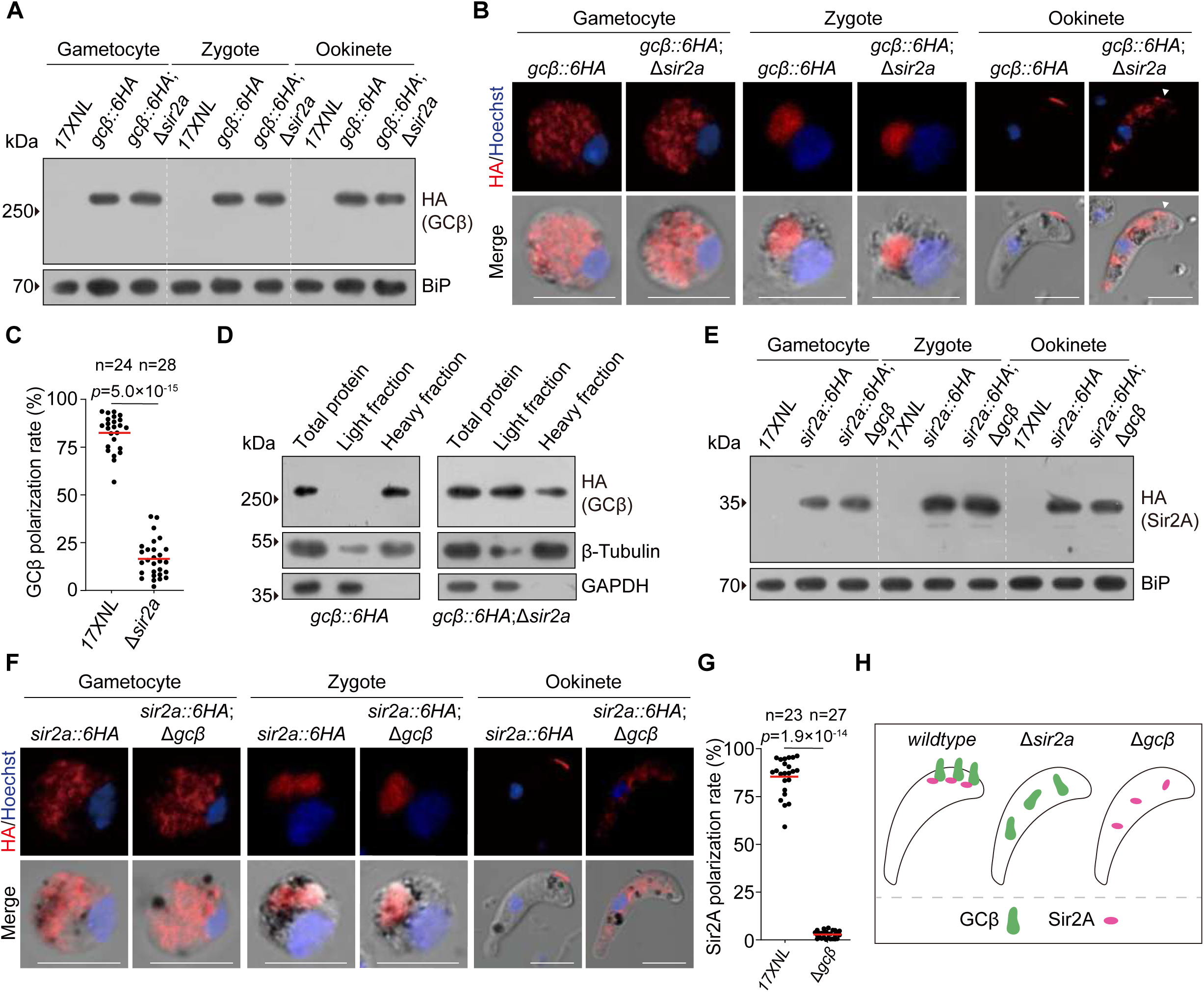
Mutual dependent localization of Sir2A and GCβ at OES in mature ookinete. **A.** Immunoblot of HA-tagged GCβ in gametocytes, zygotes, and ookinetes of the 17XNL, *gcβ::6HA* and *gcβ::6HA*;Δ*sir2a* parasites. BiP as a loading control. Three independent experiments. **B.** IFA of HA-tagged GCβ in gametocytes, zygotes and ookinetes of the 17XNL, *gcβ::6HA*, and *gcβ::6HA*;Δ*sir2a* parasites. Scale bar: 5 μm. Three independent experiments. **C.** Quantification of GCβ polarization level at OES in ookinetes in **B**. n is the number of ookinetes analyzed. Red lines show the mean value. Two-sided Mann-Whitney *U* test. **D.** Cell fractionation analysis of GCβ in *gcβ::6HA* and *gcβ::6HA*;Δ*sir2a* ookinetes via immunoblot. Light fraction includes cytosolic proteins while heavy fraction includes membrane and cytoskeleton proteins. GAPDH is a cytosolic protein and β-Tubulin is a cytoskeleton protein. Two independent experiments. **E.** Immunoblot of HA-tagged Sir2A in gametocytes, zygotes, and ookinetes of the 17XNL, *sir2a::6HA* and *sir2a::6HA*;Δ*gcβ* parasites. BiP as a loading control. Three independent experiments. **F.** IFA of HA-tagged Sir2A in gametocytes, zygotes, and ookinetes of the 17XNL, *sir2a::6HA* and *sir2a::6HA*;Δ*gcβ* parasites. Scale bar: 5 μm. Three independent experiments. **G.** Quantification of Sir2A polarization level at OES in ookinetes in **F**. n is the number of ookinetes analyzed. Red lines show the mean value. Two-sided Mann-Whitney *U* test. **H.** Cartoon showing mutual dependent localization of Sir2A and GCβ at OES in ookinete.

We next investigated whether GCβ in turn influences the Sir2A localization at OES in ookinete. We deleted the *gcβ* gene in the *sir2a::6HA* parasite and obtained the mutant clone *sir2a::6HA*;Δ*gcβ*. Deleting *gcβ* had less impact on Sir2A protein abundance in gametocytes, zygotes, and ookinetes (Figure 4E). Interestingly, GCβ depletion did not affect the cytosolic distribution of Sir2A in gametocytes and zygotes, but Sir2A lost polarization at OES and was distributed at cytosol in the *sir2a::6HA*;Δ*gcβ* ookinetes (Figure 4F and G). Therefore, Sir2A and GCβ are localized at OES in ookinetes in a mutually dependent manner (Figure 4H).

We additionally analyzed the effect of Sir2A depletion on the protein expression and localization of PDE· (cGMP-degrading enzyme) and PKG (direct effector of cGMP). The *sir2a* gene was deleted in two parasite lines *pdeδ::4Myc* and *pkg::4Myc*^19^, and we obtained two mutant lines *pdeδ::4Myc*;Δ*sir2a* and *pkg::4Myc*;Δ*sir2a*. Deleting *sir2a* had less impact on protein abundance and localization of PDE· and PKG in ookinetes (Supplementary Figure 4).

### GCβ is acetylated at gametocytes and deacetylated at mature ookinetes

Since Sir2A is a putative deacetylase, we suspected GCβ as a substrate of Sir2A. We tested whether GCβ/CDC50A is acetylated in gametocytes and deacetylated in ookinetes. Immunoblot of the HA antibody-immunoprecipitated GCβ from *gcβ::6HA* gametocyte extracts detected acetylation signals for GCβ::6HA using the pan-acetylation antibody Ac-K (Figure 5A). We generated another parasite line *gcβ::3V5* in which the endogenous GCβ was C-terminally tagged with a 3V5 and also detected the acetylation signals for GCβ::3V5 from gametocyte extracts (Figure 5B). In the parallel experiments, we analyzed the acetylation state of CDC50A and detected no acetylation signal for CDC50A::6HA from the *50a::6HA* gametocyte (Figure 5C). These results demonstrated the acetylation in GCβ but not in CDC50A in gametocytes. Furthermore, we investigated the GCβ acetylation dynamic during the gametocyte-zygote-ookinete development of the *gcβ::6HA* parasite. The acetylation signal was detected for the GCβ::6HA in gametocytes and zygotes but not in ookinetes (Figure 5D). These results indicated that GCβ is acetylated in gametocytes and zygotes but is deacetylated in mature ookinetes.

**Figure 5.**
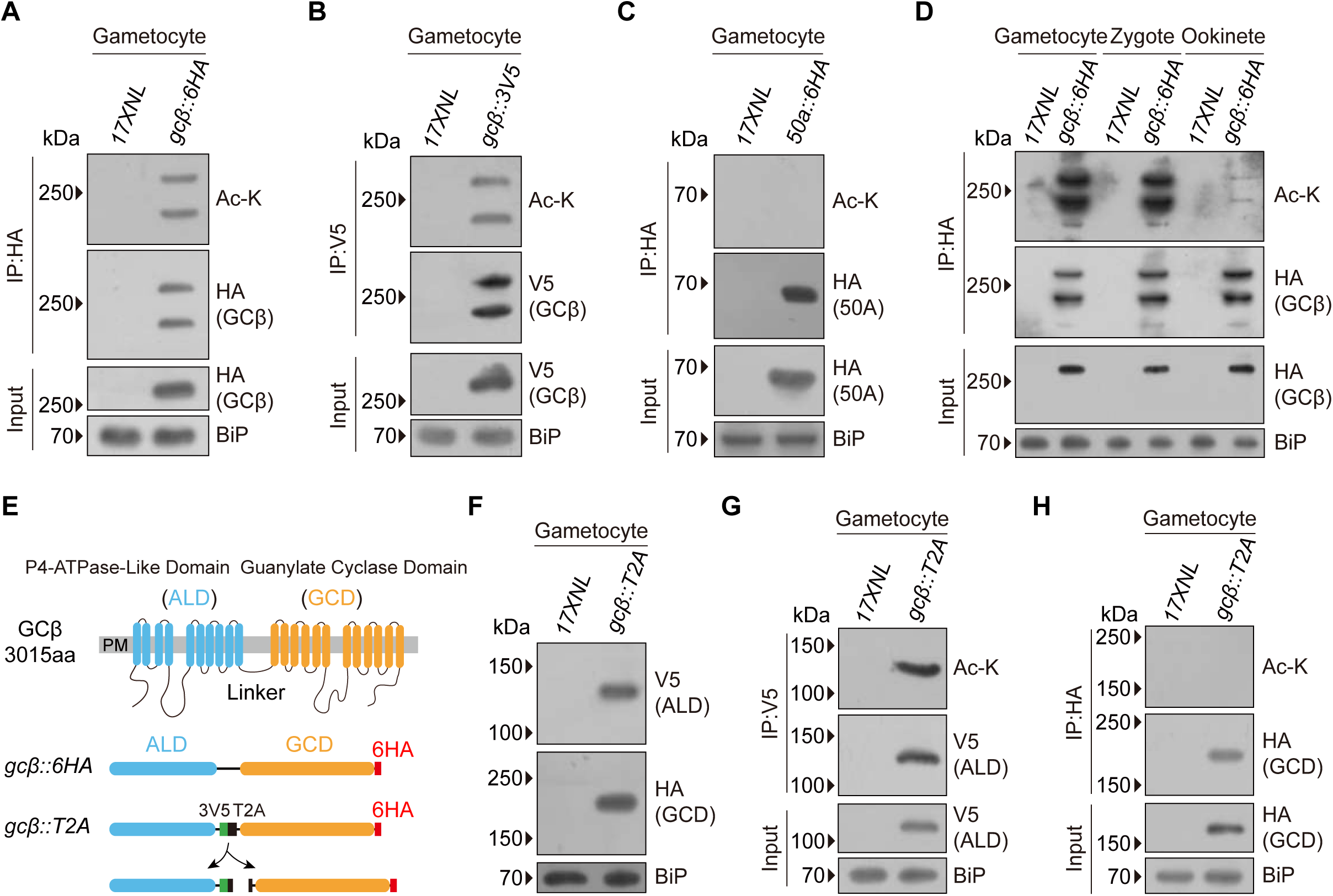
GCβ is acetylated at gametocytes and deacetylated at mature ookinetes. **A.** Detection of GCβ acetylation in 17XNL and *gcβ::6HA* gametocytes. HA-tagged GCβ was immunoprecipitated using anti-HA antibody, and the precipitates were analyzed using an anti-acetyl-lys antibody (Ac-K). BiP as a loading control. Three independent experiments. **B.** Detection of GCβ acetylation in 17XNL and *gcβ::3V5* gametocytes. V5-tagged GCβ was immunoprecipitated using anti-V5 antibody, and the precipitates were analyzed using Ac-K. Three independent experiments. **C.** Detecting of CDC50A acetylation in 17XNL and *cdc50a::6HA* gametocytes. HA-tagged CDC50A was immunoprecipitated using anti-HA antibody, and the precipitates were analyzed using Ac-K. Three independent experiments. **D.** GCβ acetylation dynamics in gametocytes, zygotes, and ookinetes of 17XNL and *gcβ::6HA* parasites. HA-tagged GCβ was immunoprecipitated using anti-HA antibody, and the precipitates were analyzed using Ac-K. Three independent experiments. **E.** Diagram of the *gcβ::T2A* line with a T2A peptide inserted into the linker region of endogenous GCβ in *gcβ::6HA* parasite. T2A allows separated expression of the V5-tagged ALD (P4-ATPase-like domain) and HA-tagged GCD (guanylate cyclase domain) of GCβ. **F.** Immunoblot confirming separated expression of V5-tagged ALD and HA-tagged GCD in *gcβ::T2A* gametocytes. Three independent experiments. **G.** Detecting of acetylation in ALD of GCβ in *gcβ::T2A* gametocytes. V5-tagged ALD was immunoprecipitated using anti-V5 antibody, and the precipitates were analyzed using Ac-K. Three independent experiments. **H.** Detecting of acetylation in GCD of GCβ in *gcβ::T2A* gametocytes. HA-tagged GCD was immunoprecipitated using anti-HA antibody, and the precipitates were analyzed using Ac-K. Three independent experiments.

The *P. yoelii* GCβ is a 3,015 aa protein that contains 22 transmembrane helixes spanning an N-terminal ALD (1-1,248 aa) and a C-terminal GCD (1,249-3,015 aa)^19^ (Figure 5E). Bioinformatic predicted 59 potential acetylation residues in GCβ using the software GPS-Palm 4.0 (https://pail.biocuckoo.org/)^36^. We attempted to map the acetylated residues in GCβ using mass spectrometry but failed to collect enough endogenous protein. Alternatively, we characterized the acetylated domain of either ALD and (or) GCD in GCβ. We used a previously generated parasite line *gcβ::T2A*^19^, in which a 88ribosome skip99 T2A peptide was introduced into the linker region between ALD and GCD in the *gcβ::6HA* parasite (Figure 5E). The T2A peptide allows separate expression of the 3V5-tagged ALD and 6HA-tagged GCD peptides, which was confirmed by immunoblot in the gametocytes (Figure 5F). Using immunoprecipitation and immunoblot, the acetylation signal was detected for the 3V5-tagged ALD but not for the 6HA-tagged GCD in the *gcβ::T2A* gametocytes (Figure 5G and H). These results demonstrated that GCβ is acetylated at the N-terminal ALD domain.

### Sir2A catalyzes the deacetylation of GCβ in mature ookinete

To prove the function of Sir2A in deacetylating GCβ, we treated the *gcβ::6HA* gametocytes with nicotinamide (NAM, inhibitor of the sirtuin deacetylases) or trichostatin A (TSA, inhibitor of the HDAC deacetylases). NAM markedly promoted GCβ acetylation, whereas TSA did not influence GCβ acetylation (Figure 6A), indicating that the deacetylase of GCβ is a member of the sirtuin family. Next, we examined GCβ acetylation alteration after loss of Sir2A during the gametocyte-zygote-ookinete development by comparing the *gcβ::6HA* and *gcβ::6HA*;Δ*sir2a* parasites. In gametocytes and zygotes, no noticeable changes in the acetylation level of GCβ were observed after loss of Sir2A (Figure 6B). However, a significant increase in GCβ acetylation was detected in ookinetes of the *gcβ::6HA*;Δ*sir2a* compared to the *gcβ::6HA* (Figure 6B). These results indicate that Sir2A is responsible for deacetylating GCβ in ookinetes.

**Figure 6.**
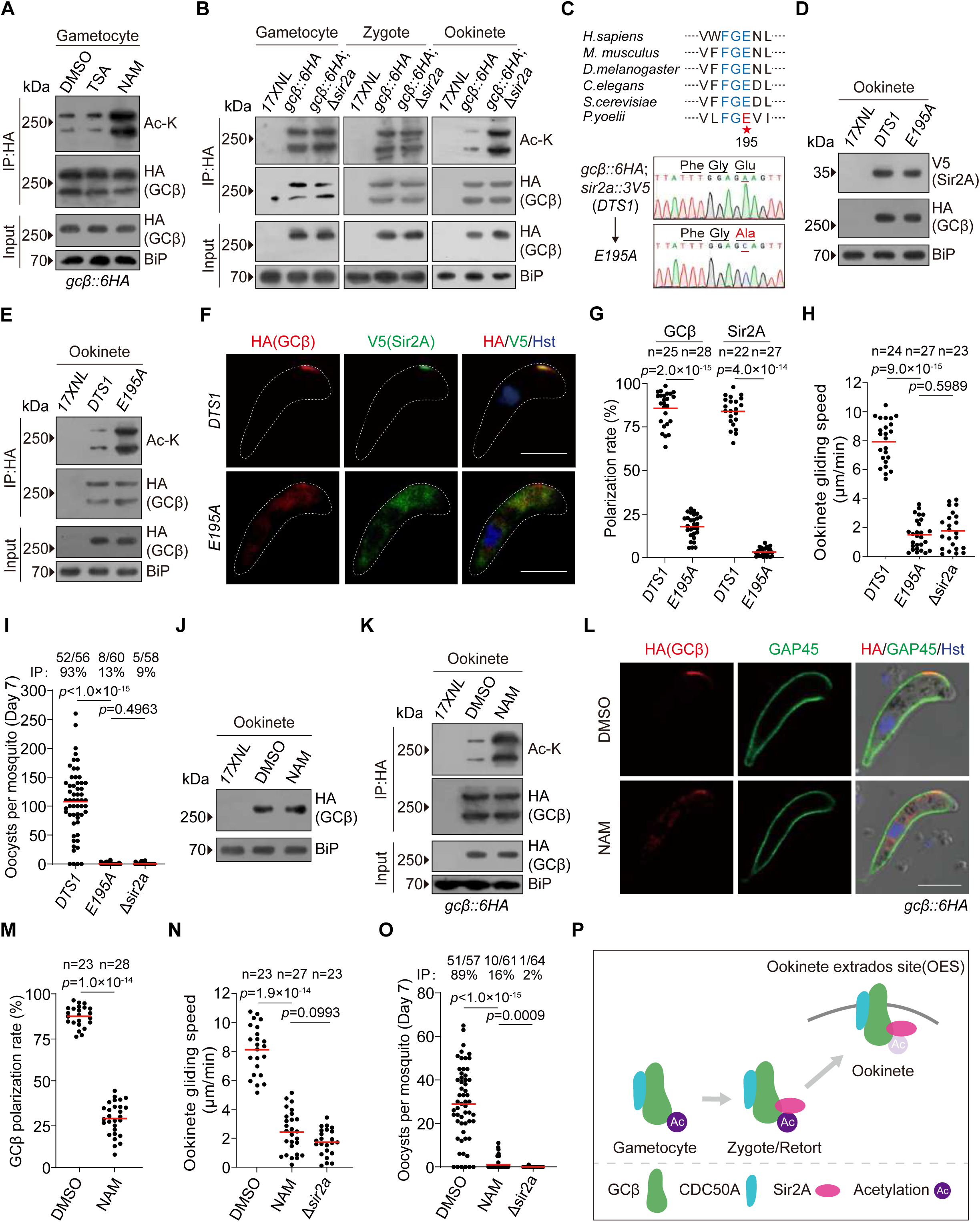
Sir2A catalyzes the deacetylation of GCβ in mature ookinete. **A.** Detection of GCβ acetylation in *gcβ::6HA* gametocytes treated with deacetylase inhibitors TSA or NAM. HA-tagged GCβ was immunoprecipitated using anti-HA antibody, and the precipitates were analyzed using Ac-K. BiP as a loading control. Three independent experiments. **B.** GCβ acetylation dynamics during the gametocyte-zygote-ookinete development of 17XNL, *gcβ::6HA*, and *gcβ::6HA*;Δ*sir2a* parasites. Three independent experiments. **C.** Generation and characterization of the mutant parasite with E195 replaced with alanine (A) in endogenous Sir2A from the parental parasite *gcβ::6HA*;*sir2a::3V5* (*DTS1*). Conserved glutamic acid (E, red star) in sirtuin proteins from several organisms is indicated. DNA sequencing confirming E195A substitution in the resulting mutant clone *E195A*. **D.** Immunoblot of HA-tagged GCβ and V5-tagged Sir2A in *DTS1* and *E195A* ookinetes. Three independent experiments. **E.** Detection of GCβ acetylation in 17XNL, *DTS1*, and *E195A* ookinetes. HA-tagged GCβ was immunoprecipitated using anti-HA antibody, and the precipitates were analyzed using Ac-K. Three independent experiments. **F.** IFA of HA-tagged GCβ and V5-tagged Sir2A in *DTS1* and *E195A* ookinetes. Scale bar: 5 μm. Two independent experiments. **G.** Quantification of GCβ and Sir2A polarization level at OES in ookinetes in **F**. n is the number of ookinetes analyzed. Red lines show the mean value. Two-sided Mann-Whitney *U* test. **H.** Ookinete gliding motility. n is the number of ookinetes analyzed. Red lines show the mean value. Two-sided Mann-Whitney *U* test. Three independent experiments. **I.** Midgut oocyst formation in mosquitoes at day 7 post infection. Red lines show the mean value, two-sided Mann-Whitney *U* test. x/y on the top is the number of mosquitoes containing oocyst/the number of mosquitoes dissected. The percentage represents the infection prevalence of mosquitoes. Two independent experiments. **J.** Immunoblot of HA-tagged GCβ in *gcβ::6HA* ookinetes after DMSO or NAM treatment. Three independent experiments. **K.** Detection of GCβ acetylation in *gcβ::6HA* ookinetes after DMSO or NAM treatment. Three independent experiments. **L.** IFA of HA-tagged GCβ in DMSO- or NAM-treated *gcβ::6HA* ookinetes. GAP45 is an IMC marker. Scale bar: 5 μm. Three independent experiments. **M.** Quantification of GCβ polarization level at OES in ookinetes in **I**. n is the number of ookinetes analyzed. Red line show the mean value. Two-sided Mann-Whitney *U* test. **N.** Ookinete gliding motility of the DMSO- or NAM-treated *gcβ::6HA* ookinetes. n is the number of ookinetes analyzed. Red line show the mean value. Two-sided Mann-Whitney *U* test. Three independent experiments. **O.** Midgut oocyst formation in mosquitos at day 7 post infection. Mosquito infection with the NAM-treated ookinetes was performed by membrane feeding using a Hemotek system. Red lines show the mean value. Two-sided Mann-Whitney *U* test. x/y on the top is the number of mosquitoes containing oocyst/the number of mosquitoes dissected. The percentage number is the mosquito infection prevalence. Three independent experiments. **P.** A model showing GCβ deacetylation by Sir2A for OES localization until ookinete maturation during ookinete development.

We verified that GCβ deacetylation depends on the deacetylase activity of Sir2A. The conserved residue Glutamic acid (E) in the salt bridge of the sirtuin proteins is critical for the binding of substrate and that E to A mutation at this residue abolishes the protein function^37^ (Figure 6C). We replaced E195 with A in Sir2A of the *gcβ::6HA*;*sir2a::3V5* (*DTS1*) parasite and generated an enzymatically inactive mutant line designated *E195A* (Figure 6C). The E195A substitution did not affect the protein level of Sir2A and GCβ in the ookinetes (Figure 6D), but resulted in significantly increased GCβ acetylation in the ookinetes of *E195A* compared to the parental parasite *DTS1* (Figure 6E). The functionally inactive protein of Sir2A-E195A lost localization at OES, causing the cytosolic distribution of the acetylated GCβ in ookinetes (Figure 6F and G). These results in the *E195A* parasites are consistent with the mutual dependent localization of Sir2A and GCβ at OES in ookinete. The *E195A* ookinetes displayed severely impaired gliding motility compared to the *DTS1* counterpart (Figure 6H) and developed no oocysts in the mosquitoes (Figure 6I), resembling the phenotype of Sir2A disruption.

We tested the effect of pharmacological inhibition of Sir2A on GCβ deacetylation and localization in ookinetes. NAM is a product of the sirtuin-mediated catalysis, blocking protein deacetylation in a negative feedback^38^. The *gcβ::6HA* zygote culture was treated with NAM at different concentrations (Supplementary Figure 6A). After 15 h treatment, NAM at 30 mΜ exerted a slight inhibition for ookinete development but influenced GCβ localization at OES in most of the developed ookinetes (Supplementary Figure 6B and C). NAM at 30 mΜ did not affect the protein abundance of GCβ (Figure 6J), but increased GCβ acetylation in the treated parasite culture (Figure 6K). GCβ lost localization at OES in the NAM-treated ookinetes (GCβ polarization rate: 87.3±6.0% in NAM, n=23; 28.3±9.5% in DMSO, n=28) (Figure 6L and M). NAM-treated ookinetes displayed severe defects in ookinete gliding (Figure 6N), resembling the defect of Δ*sir2a* in Figure 3I. Lastly, we evaluated the mosquito midgut transverse ability of the NAM-treated ookinetes. Mosquito infection with parasite was performed by membrane feeding the *in vitro* cultured ookinetes using a Hemotek system^39^. Compared to the DMSO-treated ookinetes, the NAM-treated ookinetes developed less number of oocysts in the midgut of mosquitoes on day 7 pi (Figure 6O). These results indicated that during zygote to ookinete development, Sir2A deacetylates GCβ until ookinete maturation (Figure 6P).

### Elevated NAD^+^ promotes GCβ deacetylation by Sir2A until ookinete maturation

Sir2A interacts with GCβ throughout the zygote to ookinete development but GCβ is only deacetylated by Sir2A in mature ookinete. These observations imply temporal regulation of Sir2A activity for GCβ deacetylation during the ookinete development. NAD^+^ is an essential co-substrate for sirtuin and the deacetylase activity requires a NAD^+^ level exceeding the threshold^40^. Therefore, we examined whether the level of NAD^+^ increases during the zygote to ookinete development and whether the elevated NAD^+^ activates Sir2A to deacetylate GCβ. To monitor the intracellular NAD^+^ dynamic during the zygote to ookinete development of *Plasmodium*, we used a genetically encoded NAD^+^ fluorescent biosensor FiNad^41^. This sensor binds NAD^+^ and significantly increases cpYFP fluorescence when cellular NAD^+^ levels increase^41^. FiNad was fused with mCherry for dual-color ratiometric imaging of mCherry-FiNad^41^ (Supplementary Figure 6A). mCherry-FiNad and mCherry-cpYFP (no NAD^+^ sensing as negative control) were episomally expressed in the 17XNL parasites (Supplementary Figure 6B). Both cpYFP (detecting NAD^+^) and mCherry (for expression normalization) were detected in the cytoplasm from gametocyte to ookinete for parasites expressing either mCherry-FiNad or mCherry-cpYFP (Supplementary Figure 6C). The cpYFP/mCherry ratio indicating the level of NAD^+^ did not significantly change from gametocyte to zygote in parasites expressing mCherry-FiNad. However, the ratio increased markedly during the zygote to ookinete development, reaching a maximal signal in mature ookinetes (Figure 7A and B, Supplementary Figure 6D and E). As expected, the cpYFP/mCherry ratio from mCherry-cpYFP showed no obvious fluorescence changes from gametocyte to ookinete (Figure 7A and B, Supplementary Figure 6D and E). These results indicated that the level of NAD^+^ increases during zygote to ookinete development and accumulates to its maximum in mature ookinetes.

**Figure 7.**
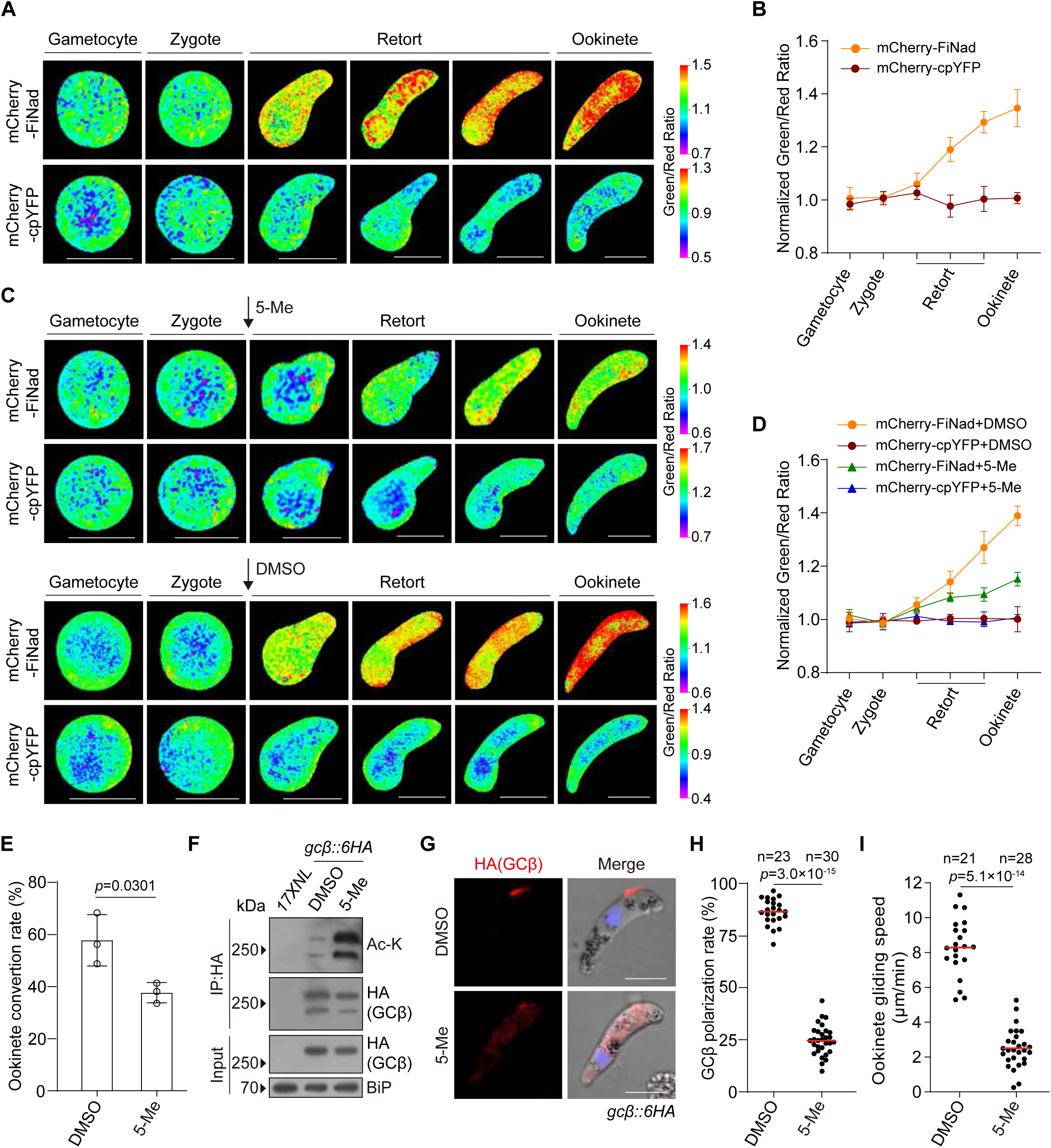
Elevated NAD^+^ promotes GCβ deacetylation by Sir2A until ookinete maturation. **A.** Detection of the NAD^+^ dynamics during gametocyte to ookinete development. The 17XNL parasites were transfected with the NAD^+^ fluorescent biosensor mCherry-FiNad (detailed information in Supplementary Figure 6). mCherry-cpYFP is a negative control sensing no NAD^+^. The cytoplasmic fluorescence of both cpYFP (488)nm for NAD^+^ detection) and mCherry (555)nm for normalization) were recorded. The ratio between cpYFP and mCherry indicates the NAD^+^ level. Scale bar: 5 μm. Three independent biological replicates. **B.** Quantification of the cpYFP/mCherry ratio in **A**. The ratio in gametocytes was set as 1.0 and all ratios in other stages were normalized. Means ± SD (n = 3 biological replicates). 30 cells were analyzed in each group of each replicate (detailed information in Supplementary Figure 6D and E). **C.** The NAD^+^ dynamics during ookinete development treated with the nicotinamidase inhibitor 5-Me-Nicotinaldehyde (5-Me). 5-Me or DMSO was added to the ookinete culture at the zygote stage (black arrow) for the parasites expressing Cherry-FiNad or Cherry-cpYFP. Scale bar: 5 μm. Three independent biological replicates. **D.** Quantification of the cpYFP/mCherry ratio in **C**. Means ± SD (n = 3 biological replicates). 30 cells were analyzed in each group of each replicate. **E.** *In vitro* mature ookinete formation of parasites treated with 5-Me or DMSO. Mean ± SD from three independent experiments, two-tailed *t*-test. **F.** Detection of GCβ acetylation in *gcβ::6HA* ookinetes treated with 5-Me or DMSO. HA-tagged GCβ was immunoprecipitated using anti-HA antibody, and the precipitates were analyzed using Ac-K. Three independent experiments. **G.** IFA of HA-tagged GCβ in *gcβ::6HA* ookinetes treated with 5-Me or DMSO. Scale bar: 5 μm. Three independent experiments. **H.** Quantification of GCβ polarization level at OES in ookinetes in **G**. n is the number of ookinetes analyzed. Red lines show the mean value. Two-sided Mann-Whitney *U* test. **I.** Ookinete gliding motility of the 5-Me- or DMSO-treated *gcβ::6HA* ookinetes. n is the number of ookinetes analyzed. Red lines show the mean value. Two-sided Mann-Whitney *U* test. Two independent experiments.

Next, we depleted the intracellular NAD^+^ to analyze its effect on GCβ deacetylation by Sir2A during the zygote to ookinete development. In the *Plasmodium*, the NAD^+^ homeostasis relies heavily on the nicotinamidase for NAD^+^ biosynthesis^42,43^. Chemical inhibition of nicotinamidase by an inhibitor 5-Me-Nicotinaldehyde (5-Me) caused significant depletion of intracellular NAD^+^ in the asexual blood stage of *P. falciparum*^43^. We tested the effect of 5-Me treatment on the level of NAD^+^ during the zygote to ookinete development in *P. yeolii*. The 17XNL zygote culture was treated with 5-Me or DMSO using a protocol similar to Supplementary Figure 5A. Compared to DMSO, 5- Me treatment inhibited the increase of cpYFP fluorescence from mCherry-FiNad during zygote to ookinete development (Figure 7C and D). In the parallel experiments, the parasites expressing mCherry-cpYFP showed no obvious fluorescence changes during ookinete development after either 5-Me or DMSO treatment (Figure 7C and D). These results indicated that 5-Me treatment decreases the NAD^+^ biosynthesis during zygote to ookinete development and depletes the NAD^+^ content in mature ookinetes. 5-Me resulted in a slight inhibition for ookinete development after 15 h treatment (Figure 7E). The drastic reduction in NAD^+^ levels after 5-Me treatment increased GCβ acetylation in the treated ookinete culture (Figure 7F) and influenced GCβ localization at OES in the developed ookinetes (Figure 7G and H). Compared to the DMSO-treated ookinetes, the 5-Me-treated ookinetes displayed defects in the ookinete gliding (Figure 7I). These results suggested that during zygote to ookinete development, the elevated level of NAD^+^ may reach the threshold to activate Sir2A for deacetylating GCβ until ookinete maturation.

## Discussion

Our previous study revealed that during the zygote to ookinete development, the GCβ/CDC50A complex translocates from cytoplasm to OES until ookinete maturation. By GCβ/CDC50A polarization at OES for elevating local cGMP concentration, mature ookinetes can activate cGMP signaling and initiate gliding motility. In this study, we identified Sir2A which forms a complex with GCβ/CDC50A during the zygote to ookinete development. Sir2A displays localization at OES in mature ookinetes similarly to GCβ/CDC50A. Notably, GCβ is acetylated in the gametocyte and maintains the acetylated status during the ookinete development. Until ookinete maturation, GCβ is deacetylated by Sir2A in the complex. Deacetylated GCβ, accompanied by CDC50A and Sir2A in the complex, translocates from the cytoplasm to OES for initiating cGMP signal. Furthermore, we revealed that the intracellular NAD^+^ increases during the zygote to ookinete development and reaches a maximum level in mature ookinete. The elevated NAD^+^ levels until ookinete maturation may induce the Sir2A-dependent GCβ deacetylation and facilitate OES translocation. We proposed a working model in Supplementary Figure 8.

The *Plasmodium* parasites encode two sirtuin proteins Sir2A and Sir2B, and each of them is conserved among different *Plasmodium* species^29^. Two previous studies successfully knocked out the *sir2a* gene in *P. falciparum*^30,31^, indicating a nonessential function for *in vitro* parasite proliferation during asexual blood stages. However, Sir2A was found to modulate the transcription of the *var* gene family by regulating the acetylation status of nuclear histones in the asexual blood stages of *P. falciparum*^30^. Whether Sir2A exerts functions in other parasite stages has not yet been investigated. In this study, we used TurboID-based proximity labeling to search the GCβ-interacting proteins in the ookinetes of the rodent malaria parasite *P. yoelii*. Among the 251 interacting proteins yielded by quantitative mass spectrometry, Sir2A is the top significant hit but Sir2B is not detected. Different from the asexual blood stage expression of Sir2A in *P. falciparum*, the expression of Sir2A is not detected or extremely low in the asexual blood stage in *P. yoelii*. The *var* gene family is unique to *P. falciparum* and no homologous of the *var* gene family exists in the genome of *P. yoelii*^44^, which is consistent with the lack of Sir2A expression in the asexual blood stage proliferation of *P. yoelii*. In mature ookinetes, Sir2A is not localized at nuclei but displayed complete concentration at OES (Figure 1D, 1F, 1H, and 2A). Sir2A forms a complex with GCβ, catalyzes GCβ deacetylation, and regulates GCβ translocation from cytoplasm to OES during the ookinete development. Besides the canonical pair of Sir2A-Histone in *P. falciparum*^30^, Sir2A-GCβ in the ookinetes represents a new sirtuin-substrate pair discovered in the *Plasmodium* parasites.

In a recent personal communication with Prof. Andy Water and Prof. Chris Janes, they generated the modified *P. berghei* parasite lines with *sir2a* gene deletion and tagging. They revealed that *P. berghei* Sir2A was co-localized with GCβ at OES in mature ookinetes and the Sir2A-knockout parasites lost ookinete gliding, resembling the defective phenotypes of the Sir2A-deficient *P. yoelii* parasite observed in this study. These results indicated that Sir2A exerts function in ookinete gliding which is conserved in *P. berghei* and *P. yoelii* parasites. In a recent reverse genetic screen in *P. berghei*, Chiamaka *et al* characterized a transmembrane protein STONES which is associated with OES and required for ookinete motility^45^. Interestingly, the homolog protein of STONES in *P. yoelii* was included in the list of proteins identified by TurboID-based proximity labeling of GCβ in this study (Supplementary Data 1). To date, the component proteins localizing at OES in ookinete include GCβ, CDC50A, Sir2A, and STONES, although more proteins may be discovered.

Acetylation and deacetylation are reversible protein post-translational modifications that regulate protein stabilization, enzymatic activity, localization, and protein-protein interaction^24^. Recently, the importance of protein acetylation and deacetylation in *Plasmodium* has been recognized, including the asexual blood stage proliferation and gametocytogenesis^46–50^. Our study demonstrates that GCβ is acetylated in the female gametocyte and maintains being acetylated during the ookinete development. Until ookinete maturation, the deacetylation of GCβ directs the protein translocation from cytoplasm to OES and initiates gliding of mature ookinetes. Two possible mechanisms may explain the effect of deacetylation in GCβ localization at OES in mature ookinete. First, a high level of acetylation in GCβ may negatively affect the trafficking of the GCβ-CDC50A-Sir2A complex from cytoplasm to OES. Second, a low level of acetylation facilitates GCβ anchoring with PPM at the OES.

The *P. yoelii* GCβ is a 3,015 aa protein containing 22 transmembrane helixes. Previous transcriptome studies detected a low transcript level of *gcβ* in female gametocytes of *P. berghei* and *P. yoelii*^51,52^. Currently, it is difficult to purify enough endogenous GCβ protein from gametocytes for mass spectrometry analysis of the acetylated lysine in GCβ. Alternatively, we used a previously generated parasite line *gcβ::T2A* in which the 3V5-tagged ALD and 6HA-tagged GCD were separately expressed in the gametocytes. In the *gcβ::T2A* gametocytes, we detected the acetylation in the 3V5-tagged ALD peptide but not in the 6HA-tagged GCD peptide, indicating that the potential acetylated lysine exists in ALD (1-1,248 aa) of GCβ. In future studies, it is important to characterize the acetylated lysine residues in ALD. GCβ is a structurally unusual protein in which the C-terminal GCD is thought to be responsible for cGMP synthesis but the N-terminal ALD is functionally obscure. GCβ is acetylated at ALD, and deacetylation of ALD plays a critical role in GCβ translocation from cytoplasm to OES in mature ookinetes. However, the separated ALD is distributed in the cytoplasm and is not targeted to OES in mature ookinetes of the *gcβ::T2A* parasites^19^. These results suggest that both the deacetylation and structural integrity of protein are required for GCβ translocation from cytoplasm to OES during ookinete development.

Sir2A interacts with its substrate protein GCβ throughout ookinete development but catalyzes GCβ deacetylation until ookinete maturation, implying temporal regulation of Sir2A activity during the ookinete development. NAD^+^ functions as a cofactor or substrate for hundreds of enzymes^53,54^ and plays important roles in many cellular processes^55,56^. The deacetylase activity of sirtuin requires an appropriate level of NAD^+^^40^. Our study revealed that the level of NAD^+^ increases during the ookinete development and reaches a maximum upon ookinete maturation. The NAD^+^ accumulating to exceed the threshold functions as a signal turning on Sir2A activity for Sir2A-dependent GCβ deacetylation. Consistent with this, in model organism lifespan-extending metabolic manipulations, such as physical exercise, caloric restriction, and time-restricted feeding, function in part by increasing NAD^+^ levels and activating sirtuins^57,58^. NAD^+^ was first discovered by regulating the metabolism in yeast^56^, and thus the links between NAD^+^ and metabolism have been widely investigated^59,60^. In *Plasmodium*, elevated NAD^+^ levels during ookinete development may prepare for enhanced oxidative metabolism requirements in the upcoming ookinete gliding. The synthesis of intracellular NAD^+^ is dictated by the de novo synthesis pathway or salvage pathway^56^. In *Plasmodium*, there is only a salvage pathway^42^. In the future, it will be interesting to elucidate whether the increase of NAD^+^ levels is caused by the increase of enzyme expression or activity in the NAD^+^ salvage pathway during ookinete development.

## Methods

### Mice and mosquitoes usage and ethics statement

The animal experiments conducted in this study were approved by the Committee for Care and Use of Laboratory Animals of Xiamen University (XMULAC20220287). Female ICR mice (5 to 6 weeks old) were obtained from the Animal Care Center of Xiamen University and used for parasite propagation, drug selection, parasite cloning, and mosquito feeding. The larvae of *Anopheles stephensi* mosquitoes (*Hor* strain) were reared at 28℃, 80% relative humidity, and a 12-hour light/12-hour dark condition in a standard insect facility. Adult mosquitoes were supplemented with 10% (w/v) sugar solution containing 0.05% 4-aminobenzoic acid and kept at 23℃.

### Plasmid construction and parasite transfection

The parasite CRISPR/Cas9 plasmid pYCm was used for gene editing^61,62^. To construct vectors for gene deletion, the 59 and 39 genomic fragments (400-800 bp) of the target gene were amplified as the left and right homologous templates respectively, and inserted into the pYCm vector. To construct vectors for gene tagging, the 59- and 39- flanking sequences (400-800 bp) at the designed insertion site of target genes were amplified as the left and right homologous templates respectively. DNA fragments encoding 6HA or 3V5 were inserted between the homologous templates in the frame with the coding sequence of the target gene. To construct vectors for nucleotide replacement, the homologous template comprises a DNA fragment spanning 582 bp upstream and 249 bp downstream of the target nucleotide in the *sir2a* gene. At least two sgRNAs were designed for each modification using the online program EuPaGDT (http://grna.ctegd.uga.edu/). Paired oligonucleotides for sgRNA were denatured at 95°C for 3 minutes, annealed at room temperature for 5 minutes, and ligated into pYCm. The sequences of primers and oligonucleotides used in this study are listed in Supplementary Table 1. The schizont-infected erythrocytes were isolated from infected mice for parasite electroporation using a 60% Nycodenz density gradient centrifugation. Parasites were electroporated with 5 μg plasmid using a Nucleofector 2b Device (Lonza, Germany). Transfected parasites were immediately intravenously injected into a naïve mouse and exposed to pyrimethamine (6 mg/mL) provided in mouse drinking water 24 hours after injection.

### Genotyping of genetically modified parasites

All genetically modified parasites (listed in Supplementary Table 2) were generated from the *P. yoelii* 17XNL or 17XNL-derived parasite lines. 10 μL parasite-infected blood samples were collected from the infected mice tail vein and lysed using 1% saponin in PBS. After centrifugation at 13,000 g for 5 minutes, the pellets were washed twice with PBS, boiled at 95°C for 10 minutes, and centrifuged at 13,000 g for 5 minutes. Supernatants containing parasite genomic DNA were subjected to genotyping. For each gene modification, the 59 and 39 homologous recombination events were detected by diagnostic PCR, confirming the successful integration of homologous templates (Supplementary Figure 8). Parasite clones with targeted modifications were obtained by limiting dilution cloning. At least two clones of each gene-modified parasite were used for phenotypic analysis. Modified parasite clones subject to additional modification were negatively selected to remove pYCm plasmid. Mice infected with pYCm plasmid-carrying parasites were exposed to 5-Fluorocytosine (Sigma-Aldrich, cat#F6627) in drinking water (2.0 mg/mL) for 6-8 days. After negative selection, two pairs of pYCm-specific primers are used to survey the residual plasmids. Clearance of plasmid in parasites after negative selection was confirmed by checking the parasite survival after reapplying pyrimethamine pressure (6 μg/mL) in mice.

### Parasite asexual blood stage proliferation in mouse

Four ICR mice were included in each group. After intravenous injection of 1.0×10^5^ parasites, parasite proliferation was monitored by Giemsa-stained thin blood smears every 2 days from day 2 to 14 post infection. The parasitemia was calculated as the ratio of parasitized erythrocytes over total erythrocytes.

### Gametocyte induction in mouse

ICR mice were treated with phenylhydrazine (80 µg/g mouse body weight; Sangon Biotech, China, cat#A600705-0025) via intraperitoneal injection. Three days post injection, the mice were infected with 5.0×10^6^ parasites through intravenous injection. Gametocytemia usually peaks at day 3 post infection. Male and female gametocytes were counted after Giemsa-stained thin blood smears. Male or female gametocytemia was calculated as a percentage of the number of male or female gametocytes over the number of parasitized erythrocytes.

### Gametocyte purification

Gametocytes were purified using the method described previously^63^. ICR mice were intraperitoneally treated with phenylhydrazine 3 days before parasite infection. From day 3 post parasite infection, the mice were orally administered 0.12 mg/day of sulfadiazine (Sigma-Aldrich, cat#S8626) for 2 days to eliminate asexual blood stage parasites. Approximately 1 mL of gametocyte-containing mouse blood was collected from the orbital sinus and suspended in 6 mL gametocyte maintenance buffer (GMB, 137 mM NaCl, 4 mM KCl, 1 mM CaCl_2_, 20 mM glucose, 20 mM HEPES, 4 mM NaHCO_3_, 0.1% BSA, and pH 7.2), the 7 mL parasite sample was layered on top of a 2 mL 48% Nycodenz solution (27.6% w/v Nycodenz in 5 mM Tris-HCl, 3 mM KCl, 0.3 mM EDTA, and pH 7.2) in a 15 mL Falcon tube. After centrifugation at 1900 g for 20 minutes, the gametocytes were collected from the interface layer and washed twice with GMB for further use.

### Male gametocyte exflagellation assay

2.5 µL of gametocyte-containing mouse blood was mixed with 100 µL of exflagellation medium. The exflagellation medium was composed of RPMI 1640 medium supplemented with 100 µM xanthurenic acid (XA, Sigma-Aldrich, cat#D120804), 2 unit/mL heparin, and pH 7.4. The mixture was incubated at 22℃ for 10 minutes. The number of parasite exflagellation centers (ECs) and total red blood cells were counted within a 1×1-mm square area of a hemocytometer under the light microscope. The exflagellation rate was calculated as the number of ECs per 100 male gametocytes.

### *In vitro* ookinete culture and purification

Mouse blood carrying 6-10% gametocytemia was collected and mixed with ookinete culture medium (RPMI 1640, 10% FCS, 100 μM XA, 25 mM HEPES, 100 μg/mL streptomycin, 100 U/mL penicillin, and pH 8.0). The culture was put at 22°C for 12-15 hours for gametogenesis, fertilization, and ookinete development. Ookinetes formation was evaluated based on cell morphology in Giemsa-stained thin blood smears. The mature ookinete conversion rate was calculated as the number of crescent-shaped mature ookinete (stage V) over that of total ookinetes (from stage I to V). Mature ookinete was purified using Nycodenz density gradient centrifugation as described previously^64^. After centrifugation at 500 g for 5 minutes, ookinete pellets were resuspended with 7 mL PBS and transferred onto the top of 2 mL of 63% Nycodenz in a 15 mL Falcon tube. After centrifuging at 1000 g for 20 minutes, the interface layer enriched with ookinetes was collected from the tube. The purity of ookinetes was examined by hemocytometer analysis. Ookinetes with more than 80% purity were used for further experiments.

### Parasite infection in mosquito

100 female *Anopheles stephensi* mosquitoes were allowed to feed on an anesthetized mouse with 4-6% gametocytaemia for 30 minutes. To evaluate midgut infection of the parasite, 30 mosquitoes were dissected and the midguts were stained with 0.1% mercurochrome 7 days post feeding. The number of oocysts in each midgut was counted under the microscope. For quantifying salivary gland sporozoites, the salivary glands were dissected from mosquitoes 14 days after feeding. Sporozoites from 30 mosquitoes were counted using a hemocytometer, and the average number of sporozoites per mosquito was calculated.

### Mosquito membrane feeding with ookinetes

1.0×10^7^ purified ookinetes from the culture were mixed with 1 mL of naïve mouse blood. The ookinete and blood mixture were added to the membrane feeder and fed to 60 female mosquitoes for 30 minutes using the Hemotek (6W1, Hemotek Limited, England). Fully engorged mosquitoes were transferred to the new container and maintained under standard conditions after feeding. 30 mosquitoes were dissected and the midguts were stained with 0.1% mercurochrome 7 days post feeding, the number of early oocysts in each midgut was counted under the microscope.

### Ookinete gliding assay

All procedures were performed in a temperature-controlled room at 22°C. 20 μL of the suspended ookinete cultures were mixed with 20 μL of Matrigel (BD Biosciences, cat#356234) on ice. The ookinete and Matrigel mixtures were transferred onto a slide, covered with a coverslip, and sealed with nail varnish. The slide was rested for 30 minutes before observation under the microscope. After tracking a gliding ookinete under the microscope, time-lapse videos (1 frame per 20 seconds, for 20 minutes) were taken to track ookinete movement using a Nikon ECLIPSE E100 microscope fitted with an ISH500 digital camera controlled by ISCapture v3.6.9.3N software (Tucsen). Ookinete motility speeds were calculated with ImageJ software using the MtrackJ plugin^65^.

### Antibodies and antiserum

The primary antibodies included: rabbit anti-HA (Cell Signaling Technology, cat#3724S, 1:1000 for immunoblot (IB), 1:1000 for immunofluorescence (IF), 1:1000 for immunoprecipitation (IP)), rabbit anti-acetylated Lys (Cell Signaling Technology, cat#9441, 1:1000 for IB), mouse anti-HA (Cell Signaling Technology, cat#2367S, 1:500 for IF), mouse anti-V5 (GenScript, A01724-100, 1:1000 for IF, 1:1000 for IB, 1:1000 for IP), mouse anti-β-tubulin II (Sigma-Aldrich, cat#T6199, 1:1000 for IF), mouse anti-β-tubulin (Sigma-Aldrich, cat#T5201, 1:1000 for IB), mouse anti-GAPDH (Servicebio, cat#GB12002, 1:1000 for IB), mouse anti-Myc (Cell Signaling Technology, cat#2276S, 1:1000 for IF, 1:1000 for IB). The secondary antibodies included: HRP-conjugated goat anti-rabbit IgG (Abcam, cat#ab6721, 1:5000 for IB), HRP-conjugated goat anti-mouse IgG (Abcam, cat#ab6789, 1:5000 for IB), Alexa 555 goat anti-rabbit IgG (ThermoFisher Scientific, cat#A21428, 1:1000 for IF), Alexa 488 goat anti-mouse IgG (ThermoFisher Scientific, cat#A11001, 1:1000 for IF), Alexa 488 conjugated streptavidin (Invitrogen, S32354, 1:1000 for IF). The anti-serums included rabbit anti-P28 (our lab, 1:1000 for IF), rabbit anti-BiP (our lab, 1:1000 for IB), rabbit anti-GAP45 (our lab, 1:1000 for IF).

### Immunofluorescence assay

Parasites fixed in 4% paraformaldehyde were transferred to a Poly-L-Lysine coated coverslip in a 24-well plate and centrifuged at 550 g for 5 minutes. Parasites were then permeabilized with 0.1% Triton X-100 solution in PBS for 10 minutes, blocked in 5% BSA solution in PBS for 60 minutes at room temperature, and incubated with the primary antibodies diluted in 5% BSA-PBS for 1 hour at room temperature. After three washes with PBS, the coverslip was incubated with fluorescent conjugated secondary antibodies for 1 hour at room temperature. Cells were stained with Hoechst 33342, mounted in 90% glycerol solution, and sealed with nail varnish. All images were acquired and processed using identical settings on Zeiss LSM 880 or LSM 980 confocal microscopes.

### Protein extraction and immunoblot

Protein extracts from the asexual blood stage parasites, gametocytes, and ookinetes were lysed in buffer A (0.1% SDS, 1mM DTT, 50 mM NaCl, 20 mM Tris-HCl, and pH 8.0) supplemented with protease inhibitor cocktail (Medchem Express, cat#HY-K0010) and PMSF (Roche, cat#10837091001). After ultrasonication, the extracts were incubated on ice for 30 minutes followed by centrifugation at 12,000 g for 10 minutes at 4°C. The supernatant was lysed in 1× Laemmli sample buffer. After SDS-PAGE separation, samples were transferred to PVDF membrane (Millipore, cat#IPVH00010). The membrane was blocked with 5% skim milk, probed with primary antibodies for 1 hour at room temperature, rinsed 3 times with 1× TBST (20 mM Tris-HCl pH 7.5, 150 mM NaCl, and 0.1% Tween20), and incubated with HRP-conjugated secondary antibodies. After three washes with TBST, the membrane was visualized with enhanced chemiluminescence detection (Advansta, cat#K12045-D10).

### Cellular fractionation

Cellular fractionation was conducted as described previously^19^. Purified ookinetes were ruptured in the hypotonic buffer (10 mM HEPES, 10 mM KCl, and pH 7.4) after passing through a 1 mL syringe needle gently ten times. The total cell lysate was centrifuged for 15 minutes at 1000 g. The supernatant (light fraction, including cytoplasm and cytosol vesicles) and the pellet (heavy fraction, including plasma membrane, IMC, and cytoskeleton) were collected and solubilized in 1× Laemmli sample buffer for 10 minutes on ice. The solubilized protein samples were analyzed using immunoblot.

### Protein immunoprecipitation

Parasites were lysed in buffer A plus (0.01% SDS, 1 mM DTT, 50 mM NaCl, 20 mM Tris-HCl, and pH 8.0) with protease inhibitor cocktail and PMSF, centrifuged at 12,000 g for 10 minutes at 4°C before collecting the supernatant solution. Rabbit anti-HA antibody was added to the protein solution and incubated at 4°C for 12 hours on a vertical mixer. After incubation, 20 μL protein A/G beads (Pierce, cat#20423) pre-balanced with buffer A plus were added and incubated for 2 hours. The beads were washed three times with buffer A plus, mixed with an equal volume of 2× Laemmli sample buffer, and oscillated on Vortex at 500 g for 5 minutes. All samples were centrifuged at 12,000 g for 5 minutes. An equal volume of supernatant from each sample was used for immunoblot.

### Detection of protein acetylation

Parasite cells were lysed in buffer A plus (0.01% SDS, 1 mM DTT, 50 mM NaCl, 20 mM Tris-HCl, and pH 8.0) containing protease inhibitor cocktail, PMSF and 30 mM NAM, incubated on ice for 30 minutes, and centrifuged at 12,000 g for 10 minutes at 4°C before collecting the supernatant solution. Rabbit anti-HA antibody was added to the protein solution and incubated at 4°C for 12 hours on a vertical mixer. After incubation, 20 μL protein A/G beads pre-balanced with buffer A plus were added and incubated for 2 hours. The beads were washed three times with buffer A plus, mixed with an equal volume of 2× Laemmli sample buffer, and oscillated at 500 g for 5 minutes. All samples were centrifuged at 12,000 g for 5 minutes. Equal volume of supernatant from each sample were used for immunoblot. Protein acetylation was analyzed using Rabbit anti-acetylated Lys antibody.

### Protein proximity labeling and biotinylated protein pull-down

1.0×10^8^ purified ookinetes from the *gcβ::TurboID::6HA* and *gcβ::T2A::TurboID::6HA* parasites were incubated with 50 µM biotin (Sigma-Aldrich, cat#B4639) at 22 °C for 3 hours. After biotinylation, the parasites were pelleted, washed three times with 1 mL ice-cold PBS to remove biotin, and lysed via ultrasonication in buffer A supplemented with protease inhibitor cocktail and PMSF. The lysate was incubated on ice for 10 minutes and centrifuged at 14,000 g at 4 °C for 10 minutes. The supernatant was mixed with 50 µL pre-balanced streptavidin sepharose (Thermal Scientific, cat#SA10004) at 4 °C overnight. The beads were washed five times with 1 mL ice-cold buffer A and washed five times with 1 mL ice-cold PBS. The washed beads were resuspended in 200 μL 100 mM TrisHCl pH 8.5 and digested with 1 µg trypsin at 37 °C overnight.

### Peptide desalting and mass spectrometry

Trifluoroacetic acid (TFA, Sigma-Aldrich, cat#T6508) was added to the trypsin-digested sample to a final concentration of 1%, and the precipitation of sodium deoxycholate was removed by centrifugation. The resulting supernatant was desalted using in-house-made StageTips that were packed with SDB-RPS (3M EMPORE, cat#2241) and conditioned with 50 μL of 100% acetonitrile (ACN, Sigma-Aldrich, cat# 34851). After loading the supernatant onto the StageTips, centrifugation was performed at 3000 g for 5 minutes. The StageTips were washed twice with 50 μL of 1% TFA/isopropyl alcohol (Sigma-Aldrich, cat# I9030) followed by a wash with 50 μL of 0.2% TFA. The peptides were eluted in glass vials (CNW Technologies, cat#A3511040) using 80% ACN/5% NH_4_OH and dried at 45 °C using a vacuum centrifuge (Eppendorf, cat#5305). The peptide samples were resolved in 2% ACN/0.1FA for LC-MS analysis. Liquid chromatography was performed on a high-pressure nano-flow chromatography system (Elute UHPLC, Bruker Daltonics). Peptides were separated on a reversed-phase column (40 cm × 75 μm i.d.) at 50 °C packed with 1.8 µm 120 Å C18 material (Welch, Shanghai, China) with a pulled emitter tip. A solution is 0.1% FA in H_2_O, and B solution is 0.1% FA in ACN. The gradient time was 60 minutes and the total run time was 75 minutes including washes and equilibration. Peptides were separated with a linear gradient from 0 to 5% B within 5 minutes, followed by an increase to 30% B within 55 minutes and further to 35% B within 5 minutes, followed by a washing step at 95% B and re-equilibration. LC was coupled online to a hybrid TIMS quadrupole time-of-flight mass spectrometer (Bruker timsTOF Pro) via a CaptiveSpray nano-electrospray ion source. We performed data-dependent data acquisition in PASEF mode with 10 PASEF scans per topN acquisition cycle. Singly charged precursors were excluded by their position in the m/z-ion mobility plane and precursors that reached a 8target value9 of 20,000 a.u. were dynamically excluded for 0.4 minutes. We used 100 milliseconds to accumulate and elute ions in the TIMS tunnel. The MS1 m/z-range was acquired from 100 to 1700, and the ion mobility range from 1.5 to 0.7 V cm^-2^. For data-independent acquisition, we adopted the isolation scheme of 25 Da × 32 windows to cover 400-1200 mz. DIA files (raw) files were input to DIA-NN (v1.8.1)^66^ FASTA files downloaded from https://www.uniprot.org (UP000072874)^67^ were added. <FASTA digest for library-free search= and <Deep learning-based spectra, RTs, and IMs prediction= were enabled. <Generate spectral library= was also enabled. <Protein inference= was set to <gene=. Other parameters were kept at the default settings. The protein groups and precursor lists were filtered at 1% FDR, using global q-values for protein groups and both global and run-specific q-values for precursors.

### Proximity ligation assay (PLA)

PLA assay was performed to detect in situ protein interaction using a commercial kit (Sigma-Aldrich, cat#DUO92008, DUO92001, DUO92005, and DUO82049). Ookinetes were fixed with 4% paraformaldehyde for 30 minutes, permeabilized with 0.1% Triton X-100 for 10 minutes at room temperature, and blocked with a blocking solution overnight at 4°C. The primary antibodies were diluted in the Duolink Antibody Diluent and incubated with ookinetes in a humidity chamber overnight at 4°C. After removing the primary antibodies, the ookinetes were rinsed twice with wash buffer A. The PLUS and MINUS PLA probes were diluted in Duolink Antibody Diluent and ookinetes were incubated in a humidity chamber for 1 hour at 37°C. Next, ookinetes were rinsed twice with wash buffer A and incubated with the ligation solution for 30 minutes at 37°C. Ookinetes were rinsed twice with wash buffer A and incubated with the amplification solution for 100 minutes at 37°C in the dark. After rinsing twice with 1× wash buffer B and one time with 0.01× wash buffer B, ookinetes were stained with Hoechst 33342 and washed twice with PBS. Images were captured and processed in a Zeiss LSM 880 confocal microscope using identical settings.

### Scanning electron microscopy

Purified ookinetes were fixed with 2.5% glutaraldehyde in 0.1 M phosphate buffer at 4℃ overnight, rinsed three times with PBS, and fixed with 1% osmium tetroxide for 2 hours. Fixed cells were dehydrated using a graded acetone series, CO_2_-dried in a critical-point drying device, and gold-coated in a sputter coater as previously^68^. The samples were imaged using a SUPRA55 SAPPHIRE Field Emission Scanning Electron Microscope.

### Episomal expression of protein

The genes encoding the NAD^+^ probe mCherry-FiNad or mCherry-cpYFP were driven by the 59-UTR (1998 bp) of the *hsp86* gene and the 39-UTR (803 bp) of the *dhfr* gene. The expressing cassette was inserted into the pL0019-derived vector with human *dhfr* for pyrimethamine selection^19^. Purified schizonts were electroporated with 10 μg plasmid DNA. Transfected parasites were immediately intravenously injected into a naïve mouse and exposed to pyrimethamine (70 μg/mL) for 5-8 days. After pyrimethamine selection, 4.0×10^6^ parasitized erythrocytes were injected intravenously into phenylhydrazine-pretreated naïve mice to induce gametocytes and were kept under pyrimethamine pressure. Mice with high gametocytaemia were used for further study.

### Measurement of intracellular NAD^+^

To measure the intracellular NAD^+^ level in the ookinete development of *Plasmodium*, we used a genetically encoded NAD^+^ fluorescent biosensor FiNad^41^. Parasites expressing the mCherry-FiNad or mCherry-cpYFP sensor proteins were collected in 200 μL PBS, washed twice with PBS, and stained with Hoechst 33342 at room temperature for 10 minutes. After centrifuging at 500 g for 3 minutes, the parasite pellets were re-suspended in 100 μL of 3% low melting agarose (Sigma-Aldrich, cat#A9414), and transferred evenly on the bottom of a 35 mm culture dish. Parasites were placed at room temperature for 15 minutes and imaged using a Zeiss LSM 880 confocal microscope. Raw data were exported to ImageJ software as 12 bit TIF for analysis. The pixel-by-pixel ratio of the 488 nm excitation image to the 555 nm excitation image in a cell was used to pseudocolor the images in HSB color space. The RGB value (255, 0, 255) represents the lowest ratio, the red (255, 0, 0) represents the highest ratio, and the color brightness is proportional to the fluorescent signals in both channels.

### Bioinformatic searches and tools

The genomic sequences of target genes were downloaded from the PlasmoDB database (http://plasmodb.org/plasmo/app/)^69^. The sgRNA of target genes was searched using the database EuPaGDT (http://grna.ctegd.uga.edu/)^70^. The amino acid sequences of protein homologs were downloaded from UniProt (https://www.uniprot.org/). The alignment of protein sequences was analyzed with MUSCLE (Version 5.1), and aligned sequences were trimmed with TrimAl (Version 1.4.1). The acetylation residues in protein were predicted using CSS-Palm 4.0 (http://csspalm.biocuckoo.org/)^36^.

### Quantification and statistical analysis

For quantifying protein polarization at OES in ookinetes, the fluorescent signals of protein were acquired using identical parameters in the microscope and analyzed by using Fiji software^71^. 30 cells were randomly chosen in each group. For quantifying the ookinete gliding speed, images were quantified using Fiji software. Statistical analysis was performed using GraphPad Prism 8.0. Details of statistical methods are described in the figure legends.

## Acknowledgments

This work was supported by the National Natural Science Foundation of China (32170427 by J.Y., 32270503 by H.C.), the Natural Science Foundation of Fujian Province (2021J01028 by J.Y.), the 111 Project sponsored by the State Bureau of Foreign Experts and Ministry of Education of China (BP2018017 by J.Y.), and the Xiamen University Double First Class Construction Project (Biology, DFC2024001 by J.Y.).

## Author contributions

Y.S. and J.Y. designed the project. Y.S., L.W., and M.J., generated the modified parasites. Y.S. and L.W., and M.J., performed phenotype analysis, protein analysis, imaging analysis, and electron microscopy analysis. Y.S. performed the bioinformatics analysis. Y.S. and J.Y. supervised the work. Y.S., H.C., and J.Y. wrote the manuscript.

